# Lipophilic compounds restore wt function of neurodevelopmental-associated KCNQ3 mutations

**DOI:** 10.1101/2024.03.13.584879

**Authors:** Michaela A. Edmond, Andy Hinojo-Perez, Mekedlawit Efrem, Yi-Chun Lin, Iqra Shams, Sebastien Hayoz, Alicia de la Cruz, Marta E. Perez Rodriguez, Maykelis Diaz-Solares, Derek M. Dykxhoorn, Yun Lyna Luo, Rene Barro-Soria

## Abstract

A major driver of neuronal hyperexcitability is dysfunction of K^+^ channels, including voltage-gated KCNQ2/3 channels. Their slow activation and deactivation kinetics produces a current that regulates membrane potential and impedes repetitive firing. Mutations in KCNQ2 and KCNQ3 lead to a wide spectrum of neurodevelopmental disorders (NDDs), ranging from benign familial neonatal seizures to severe epileptic encephalopathies and autism spectrum disorders. However, the impact of these mutations on KCNQ channel function remains poorly understood and existing treatments have unpleasant side effects. Here we use voltage clamp fluorometry and molecular dynamic simulations to investigate how R227Q and R236C, two novel NDD-causing mutations in the voltage sensor of KCNQ3, impair channel function. We show that the two mutations perturb channel gating by two distinct mechanisms: R227Q altering voltage sensor movement and R236C altering voltage sensor-to-gate coupling. Our study further shows that polyunsaturated fatty acids (PUFAs), a novel class of ion channel modulators, primarily target the voltage sensor domain in its activated conformation and yield partial and complete restoration of wt function in R227Q– and R236C-containing channels, respectively. Our results reveal the potential of PUFAs to be developed into therapies for diverse KCNQ3-based channelopathies.

## Introduction

Neuronal excitability is regulated by a wide variety of K^+^ channels, including a heterogenous population of voltage-gated K^+^ (Kv) channels. Kv channels are widely expressed in the peripheral and central neurons, where they help to regulate resting membrane potential, action potential shape and frequency, and neurotransmitter release^1^. Members of the Kv7 family of channels (Kv7.2, Kv7.3, and Kv7.5; also known as KCNQ channels^2, 3, 4^) are found in axon initial segments and Nodes of Ranvier of multiple neuronal types, where they play a critical role in generating the sub-threshold K^+^ current (M-current) that regulates excitability^5, 6^. The slow activation of KCNQ channels during action potential initiation and their lack of inactivation leads to sustained outward K^+^ currents that promote the restoration of resting membrane potential during repetitive firing. As such, these channels have the important role of providing a brake on repetitive burst firing^7^.

Impairment of M-currents due to mutation of KCNQ3 increases neuronal excitability and delays the development of complex neuronal rhythms that contribute to a variety of NDDs, including different types of epilepsy and autism spectrum disorders^8, 9, 10, 11, 12^. The phenotypic heterogeneity of KCNQ3-associated NDDs is broad, ranging from benign familial neonatal seizures with normal cognition^13^ to more severe epileptic encephalopathy with cognitive impairment^14, 15^. Among the NDD-associated KCNQ3 variants, there is an overrepresentation of mutations in the arginine residues that serve as gating charges in the voltage sensor (S4 helix), particularly residues 227, 230, and 236^9, 10, 11, 16^. However, the impact of these variants on the molecular mechanisms underlying KCNQ3 channel function remain poorly understood. Therefore, a comprehensive investigation of the mechanisms by which different variants impact channel mechanisms is required in order to understand the etiology of KCNQ3-associated NDDs.

A mechanistic understanding of KCNQ3-based channelopathies is also crucial for the development of effective therapeutic strategies. Small molecules that target neuronal KCNQ channels have been used as anticonvulsants, including pore openers such as retigabine and its derivatives^7, 17, 18, 19^, and compounds targeting the VSD such as ICA family^20, 21, 22^. However, concerns over side effects^23, 24, 25, 26, 27^ have relegated these drugs to primarily research tools. Naturally occurring compounds, including medicinal plant extracts^28^, endocannabinoids^29^, and polyunsaturated fatty acids (PUFAs)^30, 31^, have recently emerged as novel KCNQ channel regulators with differing degrees of specificity. PUFAs are particularly interesting because they modulate a variety of voltage-gated channels^32, 33^ and because their amphipathic properties confer the ability to interact with lipid-soluble and -insoluble domains of proteins. Indeed, ketogenic diets, which increase levels of PUFAs in both the blood^34^ and the brain^35^, have been previously used to treat children with epilepsy^30, 31, 36^. However, whether PUFAs can modify neuronal excitability via an effect on KCNQ channels remains unknown.

In this study, we investigated the molecular mechanisms underlying the effects of two KCNQ3 variants associated with NDDs, R227Q and R236C. We found that R227Q resulted in gain-of-function (GOF) and R236C in loss-of-function (LOF). Moreover, the two mutations perturbed channel gating by two distinct mechanisms: R227Q altering S4 movement and R236C altering S4-to-gate coupling. We also investigated whether PUFAs could diminish the effects of these mutations and identified two compounds: N-arachidonoyl amine (NAA^+^), which partially reversed the GOF mutation R227Q, and N-arachidonoyl taurine (NAT), which reversed the LOF mutation R236C. Molecular dynamics (MD) simulations suggested that PUFAs primarily target the VSD in its activated (S4 up) conformation and form a weaker interaction with the channel’s pore, in agreement with their ability to rescue mutations that disrupt S4 movement and S4-to-gate coupling. These results reveal how NDD-associated variants disrupt KCNQ3 channel function and suggest that PUFAs have potential use as efficacious, potent, and safe compounds to restore normal physiology.

## Results

### Two NDD-associated mutations in S4 have distinct biophysical properties

We set out to investigate the mechanisms by which variants in the S4 domain of KCNQ3 impact channel function to cause NDDs^9, 10, 11^. We chose two missense mutations in the voltage sensor (S4) segment of the KCNQ3 channel, R227Q and R236C (Fig. 1A), and introduced them into the KCNQ3 channel bearing the A315T mutation (hereafter referred to as KCNQ3), which enhances membrane insertion^37, 38^. KCNQ3, KCNQ3-R227Q, and KCNQ3-R236C were expressed in homotetrameric (Fig. 1) and heterotetrameric (Suppl. Fig. 1) forms in *Xenopus* oocytes and currents recorded using two-electrode voltage clamp (TEVC) electrophysiology.

**Figure 1.**
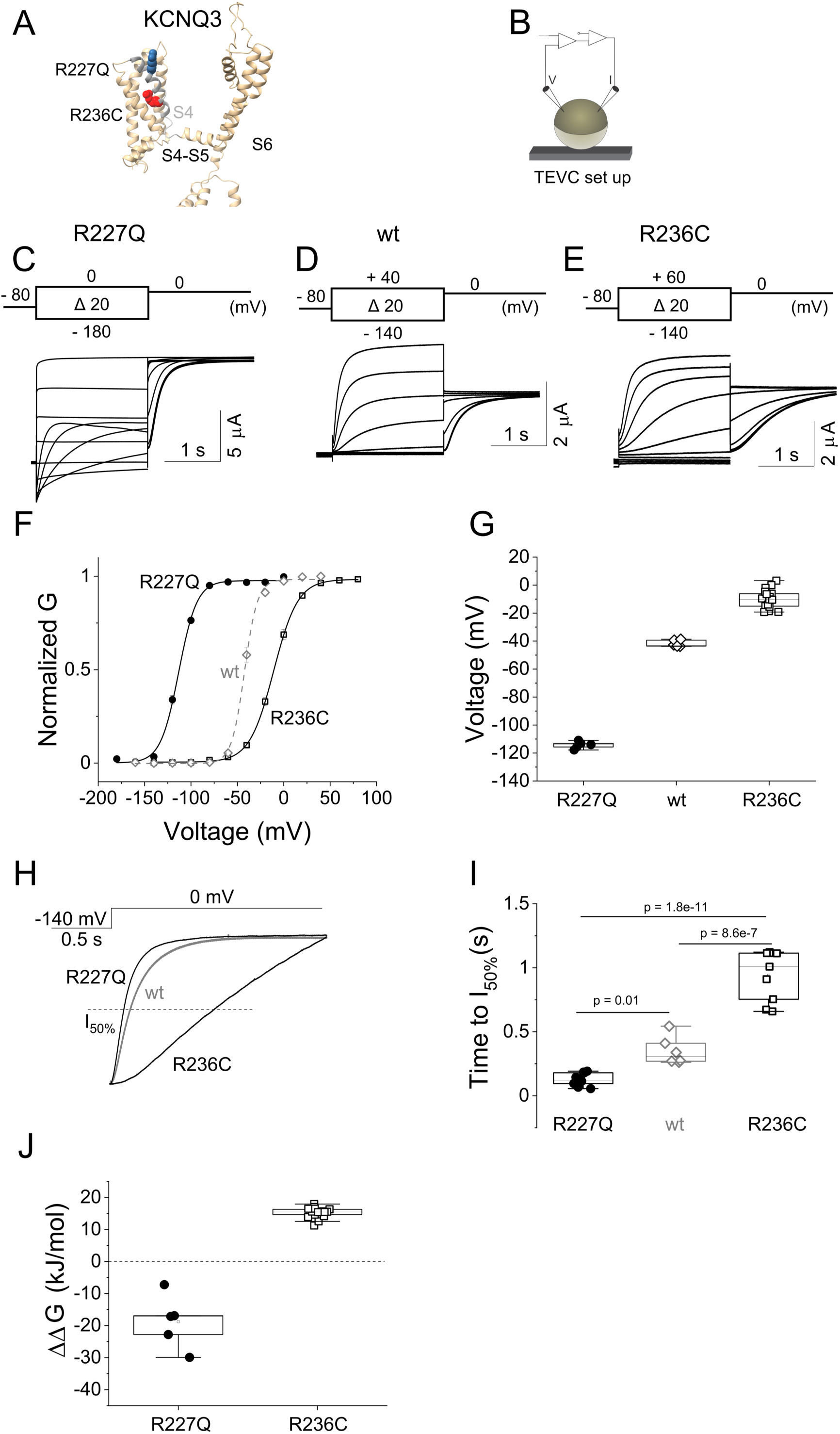
R227Q and R236C have distinct biophysical properties. (*A)* Schematic of KCNQ3 channel highlighting residues analyzed in this study. (*B*) Schematic of recording setup for TEVC. (*C-E*) Representative current traces from KCNQ3-A315T-R227Q (R227Q) (C), KCNQ3-A315T (wt) (D), and KCNQ3-A315T-R236C (R236C) (E) channels for the indicated voltage protocols. (F) Extrapolated tail conductance from panels C, D, and E were normalized and plotted against test voltages to create G(V) curves (R227Q closed circles; wt open diamonds; R236C closed squares). Lines represent the fitted theoretical voltage dependencies (eqs. 1 and 2). (*G*) Summary data for G(V) midpoints using Boltzmann fits from panel (F). (*H*) Representative current time courses of R227Q, wt, and R236C channels in response to the indicated protocol. Dashed line represents 50% of maximum current at the end of the depolarizing pulse. (*I*) Time course of current activations quantified as time to reach half maximum current at the end of the depolarizing pulse in (H, dashed line). (J) Summary of 11G data (eq. 5) for R227Q and R236C channels. Midpoints of voltage activation and 11G values for each channel are shown in Table 1. Data presented as mean ± SEM, n = 5–11. Statistical significance determined using one-way ANOVA and Bonferroni’s post hoc test, p < 0.05.

**Table 1.** Effect of NDD-associated mutations on KCNQ3 channel function. Summary data for half-activation voltage (V_1/2_ and F_1/2_) for each G(V) and F(V) relationships, shifts in half-activation voltage (ΔV_1/2_, ΔF_1/2_), Gibbs free energy associated with channel opening (11G_G_) and S4 activation (11G_F_), and relative change in potassium current at –60 mV (1I_–60_) for ‘wt’ KCNQ3 (KCNQ3-A315T), Alexa-488–5 maleimide labelled– KCNQ3-A315T-Q218C (Q218C), and unlabeled (R227Q and R236C) and labelled (Q218C/R227Q and Q218C/R236C) NDD-associated mutations. κ is the slope of the activation curve fitted with the Boltzmann equation 1 (see methods). Values are means ± S.E. n represents the number of cells analyzed. Statistical significance determined using one-way ANOVA and Bonferroni’s post hoc test, p < 0.05 vs. the corresponding value in: *wt KCNQ3 and †labeled-KCNQ3-Q218C.

| | n | $\kappa$ (g)<br>(mV) | $V_{1/2}$<br>(mV) | $\kappa$ (f)<br>(mV) | $F_{1/2}$<br>(mV) | $\Delta V_{1/2}$<br>(mV) | $\Delta F_{1/2}$<br>(mV) | $\Delta\Delta G_G$<br>(kJ/mol) | $\Delta\Delta G_F$<br>(kJ/mol) | $T_{50\%}$ I-act<br>(S) |
| --- | --- | --- | --- | --- | --- | --- | --- | --- | --- | --- |
| 'wt' KCNQ3 | 9 | $7.1 \pm 0.6$ | $-41.3 \pm 0.5$ | - | - | - | - | - | - | $0.35 \pm 0.11$ |
| R227Q | 5 | $8.5 \pm 0.7$ | $-116.9^* \pm 0.8$ | - | - | $-75.6 \pm 1.4$ | - | $-18.5 \pm 3.7$ | - | $0.13^* \pm 0.04$ |
| R236C | 11 | $13.7 \pm 0.2$ | $-8.4^* \pm 0.2$ | - | - | $+32.9 \pm 1.5$ | - | $+15.2 \pm 0.4$ | - | $0.94^* \pm 0.3$ |
| Q218C | 13 | $12.0 \pm 0.9$ | $-34.9 \pm 0.9$ | $13.9 \pm 0.6$ | $-33.9 \pm 0.7$ | $+6.4 \pm 1.4$ | - | - | - | - |
| Q218C/R227Q | 6 | $12.8 \pm 0.8$ | $-122.8^\dagger \pm 1.0$ | $12.2 \pm 0.5$ | $-127.4^\dagger \pm 0.6$ | $-87.9 \pm 0.2$ | $-94.5 \pm 1.6$ | $-9.5 \pm 1.3$ | $-11.6 \pm 2.2$ | - |
| Q218C/R236C | 5 | $14.8 \pm 0.3$ | $+6.3^\dagger \pm 0.4$ | $22.0 \pm 0.7$ | $-17.7^\dagger \pm 0.8$ | $+41.0 \pm 0.7$ | $-43 \pm 1.1$ | $+14.6 \pm 1.3$ | $+11.5 \pm 0.6$ | - |

In comparison to homomeric KCNQ3 channels, we found that homomeric R227Q channels displayed a hyperpolarizing (leftward) shift in their steady-state conductance-voltage G(V) curve and an accelerated time course of current activation (Fig. 1C,F-I, Table 1), in agreement with previous reports^9^. Indeed, homomeric R227Q channels remained open throughout the physiological voltage range (from –100 to 0 mV, Fig. 1F). In contrast, homomeric R236C channels displayed a depolarizing (rightward) shift in their G(V) curve and a slower time course of current activation (Fig. 1E-I, Table 1). To better compare the functional effect of G(V) shifts on these mutants we corrected for the differences in slopes and calculated the change in Gibbs free energy (ΔΔG,) to estimate the energy required to open each channel^39, 40^. Compared to homomeric KCNQ3 channels, R227Q decreased the energy required for channel opening whereas R236C increased it (Fig. 1J, Table 1), consistent with GOF and LOF, respectively.

Because most patients bear mutated alleles heterozygously, and the M-current primarily comprises heteromeric configurations of KCNQ2/3^41^ (the contribution of KCNQ5 to the M-current will not be considered in this study for simplicity), we co-expressed mutant-bearing KCNQ2/3 subunits heteromerically in *Xenopus* oocytes (Suppl. Fig. 1). Compared to wt heteromeric KCNQ2/3 channels (expressed in a 1:1 ratio), channels assembled from KCNQ2, KCNQ3, and KCNQ3-R227Q (wtQ2/wtQ3/Q3-R227Q in a 2:1:1 ratio to mimic heteromeric channels in a heterozygous patient case) displayed a leftward shift in the G(V) relationship (Suppl. Fig. 1A top, D-E, and Suppl. Table 1). Incorporation of a second KCNQ3-R227Q subunit into the tetramer (wtQ2/Q3-R227Q in a 1:1 ratio to mimic heteromeric channels in a homozygous patient) caused a further leftward shift in the G(V) curve (Suppl. Fig. 1A bottom, D-E, and Suppl. Table 1), revealing a gene-dose effect for this mutation. Moreover, the presence of either one or two R227Q subunits in the tetramer decreased the change in Gibbs free energy and thus the energy required to open the channel (Suppl. Fig. 1F and Suppl. Table 1). In contrast, oocytes expressing channels incorporating R236C subunits (wtQ2/wtQ3/Q3-R236C in a 2:1:1 ratio or wtQ2/Q3-R236C in a 1:1 ratio) exhibited a rightward shift in the G(V) relationship and required more free energy to open the channel (Suppl. Fig. 1C-F and Suppl. Table 1). Similar to R227Q, we found a gene-dose effect for R236C-bearing mutations, such that the biophysical phenotype was more severe with a greater number of mutated subunits per channel. This indicates that wt subunits can partially restore heteromeric KCNQ channel function. To avoid any confounding effects of subunit composition, we proceeded to investigate the mechanisms underlying GOF and LOF in these two mutants using homomeric KCNQ3 channels.

### R227Q and R236C impair channel function by different mechanisms

Given the effect of R227Q and R236C on voltage gating, we measured S4 movement and channel opening simultaneously using voltage clamp fluorometry (VCF). Wt and variant-bearing KCNQ3 channels were fluorescently labelled by attaching Alexa-488-maleimide to the Q218C site in the S3-S4 loop (hereafter referred to as KCNQ3^L^) (Fig. 2A). This manipulation has previously been shown to faithfully report S4 movement^42, 43^. The KCNQ3^L^ construct produced a robust voltage-dependent fluorescence change, F(V), that was correlated with the G(V) curve (Suppl. Fig. 2). Interestingly, KCNQ^L^-R227Q channels also exhibited G(V) and F(V) signals that closely followed each other, but were left shifted compared to KCNQ3^L^ (Fig. 2B, C and Table 1). In contrast, the G(V) and F(V) curves of KCNQ^L^-R236C channels were right shifted compared to KCNQ3^L^, although the F(V) curve showed an intermediate shift that did not overlap with the G(V) curve (Fig. 2D, E and Table 1). The changes in Gibbs free energy associated with S4 activation and channel opening in R227Q– and R236C–containing channels were consistent with their GOF and LOF phenotypes, respectively (Table 1). Together, these results suggest that R227Q and R236C shift G(V) curves by interfering with different gating transitions. The alignment of F(V) and G(V) curves at hyperpolarized voltages for the R227Q variant, together with the decrease in free energy required to activate S4, suggest that R227Q directly affects S4 movement. In contrast, the distinct F(V) and G(V) curves for the R236C variant, and increased energy required to both move S4 and open the channel, is indicative of an alteration in S4-to-gate coupling.

**Figure 2.**
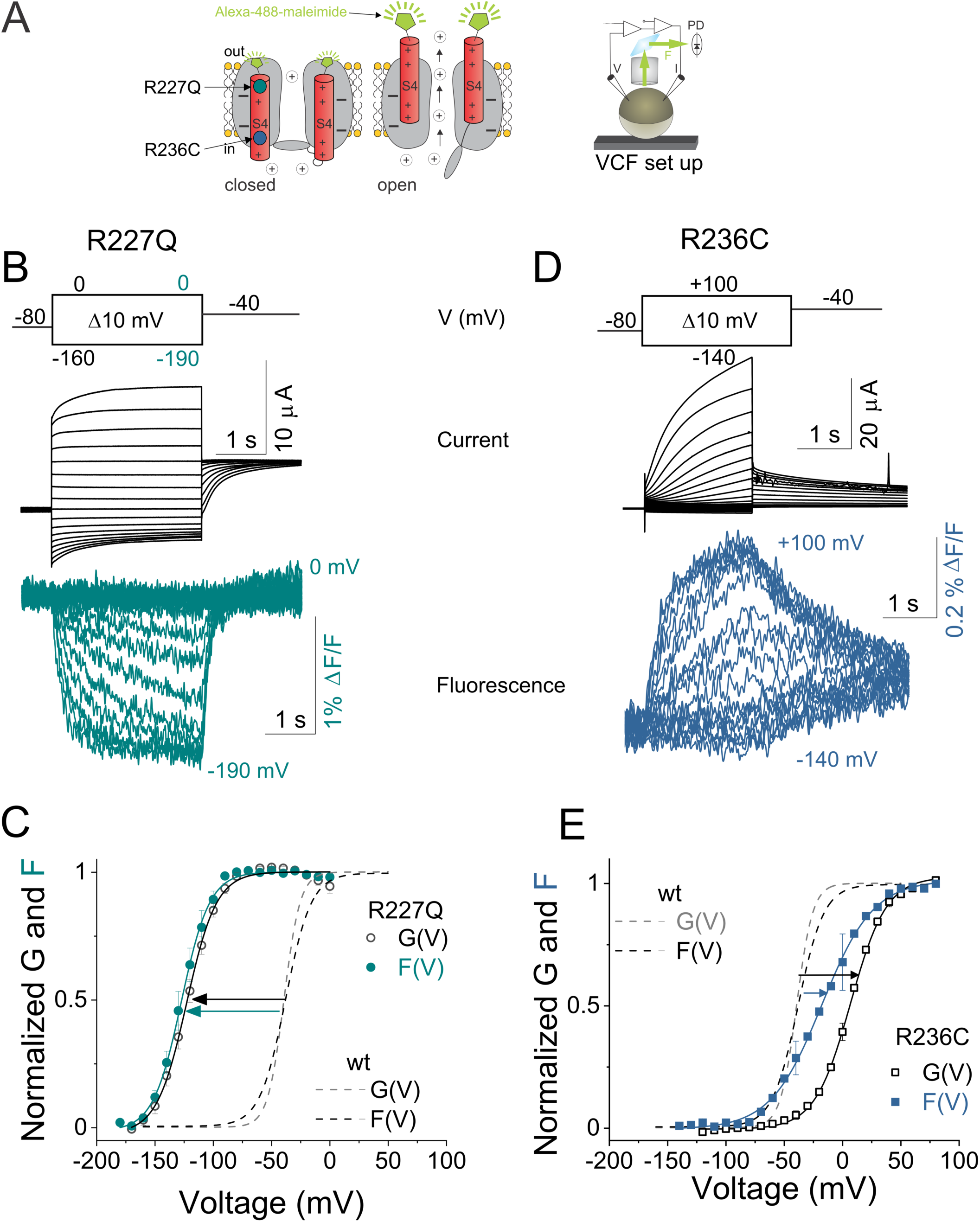
R227Q and R236C impair channel function by different mechanisms. (*A*) Schematic representing VCF technique. A cysteine introduced at position 218 (close to S4) is labeled with a fluorophore tethered to Alexa-488–5 maleimide. Upon voltage changes, labeled-S4s move and the environment around the tethered fluorophore changes, altering fluorescence intensity. Current and fluorescence are recorded simultaneously using the setup shown on the right. Location of the two NDD–associated mutations (R227Q and R2236C) are shown on the left. (*B*) Representative current (black) and fluorescence (cyan) traces from Alexa-488–labeled KCNQ3-A315T-R227Q (R227Q) channels for the indicated voltage protocol. (*C*) Normalized G(V) (black circles and black solid line from a Boltzmann fit) and F(V) (cyan circles and cyan solid line from a Boltzmann fit) curves from labeled R227Q. (*D*) Representative current (black) and fluorescence (blue) traces from Alexa-488– labeled KCNQ3-A315T-R236C (R236C) channels for the indicated voltage protocol. (*E*) Normalized G(V) (black squares and black solid line from a Boltzmann fit) and F(V) (blue squares and blue solid line from a Boltzmann fit) curves from labeled R236C. Dashed lines represent labeled pseudo-wt KCNQ3-A315T-Q218C G(V) (gray) and F(V) (black) curves for comparison (raw data shown in Table 1 and Supplementary Figure 2). Midpoints of voltage activation are shown in Table 1. Data represent mean ± SEM. n = 5–13

### PUFAs can activate or inhibit KCNQ3 channel function

Because the R227Q and R236C variants alter S4 movement and gate opening via different mechanisms, we hypothesized that different strategies would be required to restore channel functionality. We therefore screened a diverse panel of naturally occurring PUFAs and PUFA analogs for their ability to modulate G(V) relationships, current amplitudes (Gmax), and kinetics of activation and deactivation in KCNQ3 channels expressed in *Xenopus* oocytes (Fig. 3 and Suppl. Fig. 3). Extracellular application of 25 μM μ-3 docosahexaenoic acid (DHA), linoleic acid (LA), μ-6 arachidonic acid (AA), and arachidonoyl ethanolamine (O-AEA) had minimal effects on G(V) curves and only weakly increased Gmax (Suppl. Fig. 3A-H, O, P, Suppl. Table 1). In addition, measurements of Gibbs free energy (ΔΔG) revealed only small effects on the energy required to open KCNQ3 channels (Suppl. Fig. 3Q, Suppl. Table 1). We also quantified the amount of potassium current at – 60 mV (I–_60_), near the action potential threshold for most neurons, and found only minimal changes with DHA, LA, AA, and O-AEA (Suppl. Fig. 3R and Suppl. Table 1).

**Figure 3.**
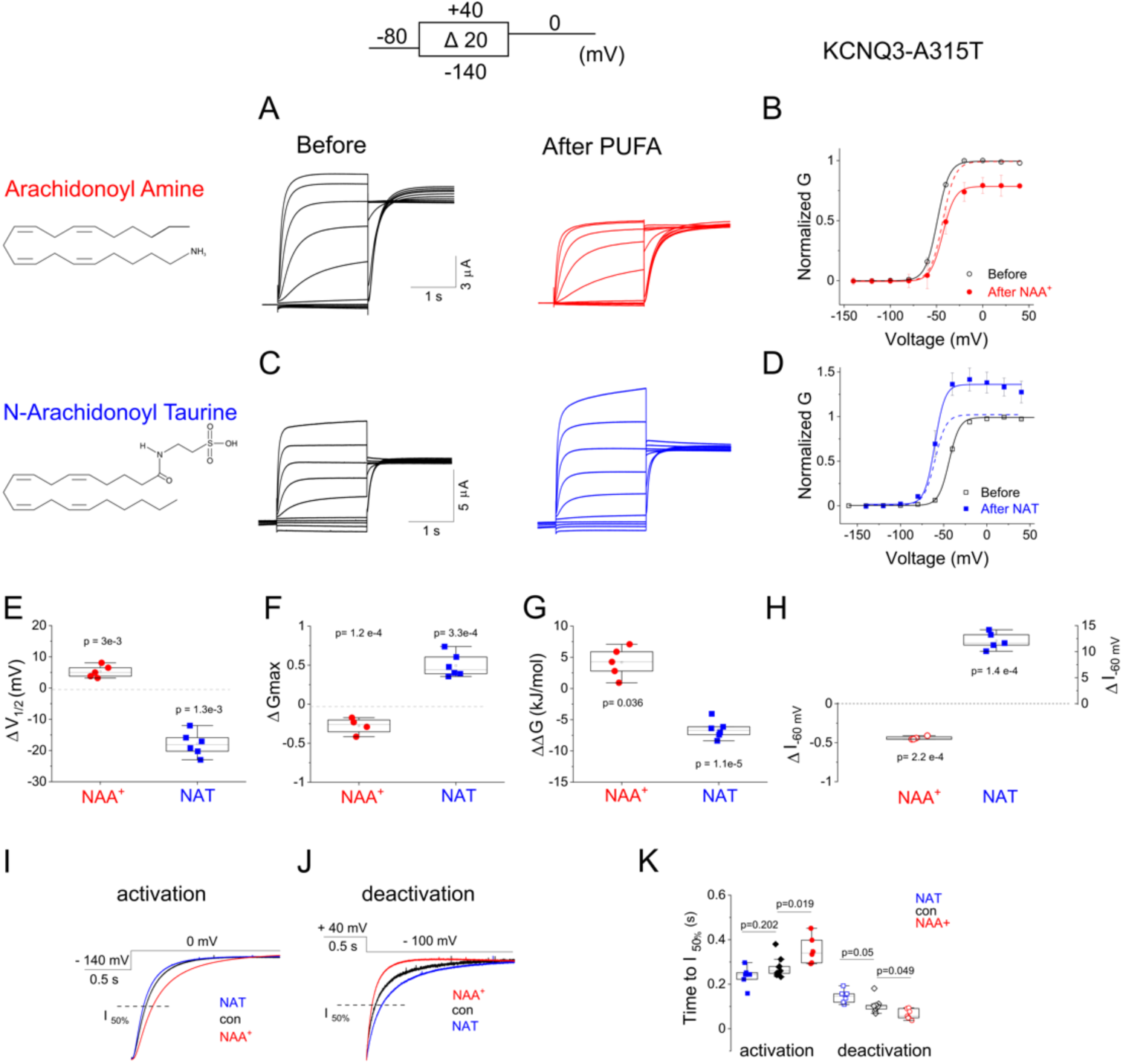
PUFAs can activate or inhibit KCNQ3 channel function. (*A and C*) Representative current traces from KCNQ3-A315T channels in the absence (before) or presence (after) of 25 μM NAA^+^ (A) or NAT (C) for the indicated voltage protocol. (*B and D*) Steady-state G(V) curves (solid lines from a Boltzmann fit) obtained from recordings in panels *A* and *C* normalized to peak conductance before PUFA application (open symbols). G(V) relationships (solid lines from a Boltzmann fit) in the presence of PUFAs shown as closed symbols and dashed lines represent their respective normalized G(V). (*E*-*H*) Summary data for shifts in half-activation voltage (ΔV_1/2_) for each G(V) (E), increase in Gmax (ΔGmax) (F), 11G (G), and relative change in potassium current at –60 mV (1I_–60_) (H) induced by 25 μM PUFAs on KCNQ3 channels. Dashed lines represent DMSO-induced changes for comparison. (*I*-*J*) Representative time courses of current activation (I) and deactivation (J) in the absence (black) and presence of NAA^+^ (red) and NAT (blue) in KCNQ3-A315T channels in response to the indicated voltage protocol. Dashed lines represent 50% maximum current level at the end of the depolarizing pulse. (*K*) Time courses of current activation (closed symbols) and deactivation (open symbols) in the absence (black) and presence of NAA^+^ (red) and NAT (blue) quantified as time to reach half maximum current level at the end of the depolarizing pulse in (I and J, dashed lines). Values for V_1/2_, ΔV_1/2_, Gmax (eq 3), 11G (eq 6), 1I_-60_ (eq 4), and time to I_50%_ given in Table 2. Data represent mean ± SEM; n = 6–9. Statistical significance determined using Student’s T-test to compare the PUFA–induced change in ΔV_1/2_, ΔGmax, 11G, 1I_–60_, and time to I_50%_ relative to mock application of a solution containing only vehicle (DMSO); only significant differences are shown. One-way ANOVA and Bonferroni’s post hoc test, containing the other 5 screened PUFAs and PUFA analogs in Suppl. Figure 3, also gave p < 0.05 for NAA^+^ and NAT (see Suppl. Figure 3).

**Table 2.** Effect of PUFAs on ‘wt’ and NDD-associated mutations of KCNQ3 channel. Summary data for half-activation voltage (V_1/2_ and F_1/2_) for each G(V) and F(V) relationships, shifts in half-activation voltage (ΔV_1/2_, ΔF_1/2_), Gibbs free energy associated with channel opening (11G_G_) and S4 activation (11G_F_), maximal current amplitude (ΔGmax), relative change in potassium current at – 60 mV (1I_–60_), and time courses of current and fluorescence activation and deactivation induced by 25 μM PUFAs on ‘wt’ KCNQ3 (KCNQ3-A315T) and mutations used in this study. Q218C represents Alexa-488–5 maleimide labelled– KCNQ3-A315T-Q218C. 5,8,11,14-cis-N-arachidonoyl amine (NAA^+^), 5,8,11,14-all-cis-N-arachidonoyl taurine (NAT) Values are means ± S.E. n represents the number of cells analyzed. Statistical significance determined using Student’s T-test or one-way ANOVA and Bonferroni’s post hoc test to compare the PUFA–induced change in ΔV_1/2_, ΔGmax, 11G, 1I_–60_, and time to I_50%_. p < 0.05 vs. the corresponding value in: *wt KCNQ3 ± DMSO, ‡wt KCNQ3 ± PUFA, and †labeled KCNQ3-Q218C ± PUFA

| | n | $V_{1/2}$<br>(mV)<br>before | $V_{1/2}$<br>(mV)<br>after PUFA | $F_{1/2}$<br>(mV)<br>before | $F_{1/2}$<br>(mV)<br>after PUFA | $\Delta V_{1/2}$<br>(mV) | $\Delta F_{1/2}$<br>(mV) | $\Delta\Delta G_G$<br>(kJ/mol) | $\Delta\Delta G_F$<br>(kJ/mol) | $\Delta G_{max}$ | $\Delta I_{60}$ | $T_{50\%}$ I-act<br>(S)<br>before | $T_{50\%}$ I-act<br>(S)<br>after PUFA | $T_{50\%}$ I-deact<br>(S)<br>before | $T_{50\%}$ I-deact<br>(S)<br>after PUFA | $T_{50\%}$ F-act<br>(S)<br>before PUFA | $T_{50\%}$ F-act<br>(S)<br>after PUFA | $T_{50\%}$ F-deact<br>(S)<br>before | $T_{50\%}$ F-deact<br>(S)<br>after PUFA |
| --- | --- | --- | --- | --- | --- | --- | --- | --- | --- | --- | --- | --- | --- | --- | --- | --- | --- | --- | --- |
| ‘wt’ KCNQ3 ± DMSO (0.03%) | 9 | -41.3 ± 0.5 | -43.0 ± 2.0 | - | - | -1.4 ± 1.5 | - | - | - | -0.05 ± 0.12 | - | - | - | - | - | - | - | - | - |
| ‘wt’ KCNQ3 ± NAA<sup>+</sup> | 6 | -43.5 ± 0.3 | -38.2* ± 0.4 | - | - | +5.3 ± 2.4 | - | +3.1 ± 1.1 | - | -0.28* ± 0.14 | -0.44* ± 0.22 | 0.28 ± 0.1 | 0.22* ± 0.1 | 0.11 ± 0.04 | 0.07* ± 0.03 | - | - | - | - |
| ‘wt’ KCNQ3 ± NAT<sup>-</sup> | 11 | -41.3 ± 0.7 | -60.2* ± 0.6 | - | - | -18.9 ± 7.7 | - | -4.9 ± 0.6 | - | +0.49* ± 0.7 | +12.8* ± 5.4 | 0.29 ± 0.1 | 0.36* ± 0.16 | 0.11 ± 0.04 | 0.15* ± 0.07 | - | - | - | - |
| R227Q ± NAA<sup>+</sup> | 7 | -118.7 ± 1.6 | -107.9† ± 1.0 | - | - | +10.8 ± 2.9 | - | +3.1 ± 0.9 | - | -0.31† ± 0.17 | -0.57† ± 0.23 | 0.12 ± 0.04 | 0.14 ± 0.4 | 0.17 ± 0.08 | 0.095† ± 0.04 | - | - | - | - |
| R236C ± NAT<sup>-</sup> | 9 | -5.8 ± 1.4 | -24.1† ± 4.7 | - | - | -18.3 ± 4.4 | - | -4.2 ± 0.7 | - | -0.44† ± 0.16 | +7.5† ± 3.1 | 0.72 ± 0.3 | 0.43† ± 0.16 | 0.23 ± 0.09 | 0.36† ± 0.15 | - | - | - | - |
| R227Q ± NAT<sup>-</sup> | 5 | -116.4 ± 0.8 | -119.3 ± 0.5 | - | - | -2.9 ± 1.2 | - | -0.74 ± 0.4 | - | +0.59† ± 0.26 | - | - | - | - | - | - | - | - | - |
| K146Q ± NAT<sup>-</sup> | 8 | -19.3 ± 3.9 | -26.8† ± 5.2 | - | - | -7.5 ± 2.8 | - | -1.1 ± 0.3 | - | +0.21 ± 0.09 | - | - | - | - | - | - | - | - | - |
| K284Q ± NAT<sup>-</sup> | 6 | -40.7 ± 1.0 | -49.3† ± 1.5 | - | - | -8.6 ± 3.5 | - | -2.4 ± 1.1 | - | +0.18 ± 0.07 | - | - | - | - | - | - | - | - | - |
| Q218C/R227Q ± NAA<sup>+</sup> | 9 | -122.8 ± 0.3 | -115.6† ± 0.7 | -127.5 ± 0.9 | -122.0 ± 1.1 | +7.2 ± 4.3 | +5.5 ± 1.4 | +3.7 ± 2.5 | +3.3 ± 2.2 | -0.37† ± 0.09 | -0.7† ± 0.3 | 0.18 ± 0.001 | 0.25 ± 0.002 | 1.8 ± 0.8 | 1.4 ± 1.1 | 0.1 ± 0.06 | 0.15 ± 0.09 | 0.33 ± 0.08 | 0.22† ± 0.07 |
| Q218C/R236C ± NAT<sup>-</sup> | 5 | +6.3 ± 0.4 | -2.8† ± 2.4 | -17.7 ± 0.8 | -26.1 ± 3.4 | -9.1 ± 1.3 | -8.4 ± 2.1 | -4.8 ± 0.9 | -4.0 ± 2.0 | +0.23† ± 0.07 | +22.3† ± 7.3 | 1.7 ± 0.003 | 0.77† ± 0.002 | 2.0 ± 0.9 | 2.3 ± 1.0 | 0.39 ± 0.22 | 0.22† ± 0.13 | 0.77 ± 0.44 | 0.97 ± 0.56 |

However, two PUFA analogs, N-arachidonoyl amine (NAA^+^) and N-arachidonoyl taurine (NAT), had more substantive effects on KCNQ3 channels. 50 μM NAA^+^ right shifted the G(V) relationship by ∼5 mV, reduced Gmax by 27%, increased the energy required for channel opening by ∼ + 4.2 kJ/mol, and reduced I–_60_ by 40% (Fig. 3A-B, E-H, and Table 2). NAA^+^ also slowed the time course of current activation and accelerated the time course of current deactivation (Fig. 3I-K). In contrast, 25 μM NAT activated KCNQ3 channels by left shifting the G(V) relationship by ∼19 mV, increasing Gmax, decreasing the energy required for channel opening by ∼7 kJ/mol, and increasing I–_60_ by 12-fold (Fig. 3C-K and Table 2). In addition, NAT slightly accelerated the time course of current activation and slowed the time course of current deactivation (Fig. 3I-K and Table 2). Thus, although both NAA^+^ and NAT modulate KCNQ3 channels, they act via mechanisms that inhibit and activate channel function, respectively.

### PUFAs primarily target the KCNQ3 voltage sensor

Given its activating effect, we wondered whether NAT might activate KCNQ3 channels by interacting with the proposed binding site for the KCNQ channel openers retigabine^44^ and ICA-069673^45^. These compounds bind to W265 (in the S5 helix), and P211 and L198 (in the voltage-sensing domain) (Suppl. Fig. 4A). We therefore tested the effect of NAT on KCNQ3-W265L, KCNQ3-P211N, and KCNQ3-L198F channels, and observed similar activation to that seen in wt KCNQ3 channels (Suppl. Fig. 4A and Suppl. Table 1). These data suggest that the activating effects of NAT are not dependent on W265, P211, or L198 and therefore that NAT activates KCNQ3 channels via a distinct mechanism.

To identify the sites at which NAT binds to KCNQ3 channels, we conducted all-atom MD simulations of two KCNQ3 homology models in the active (S4 up) and resting (S4 down) conformation in a solvated POPC bilayer. We incorporated 20 NAT molecules in the upper leaflet and 23 PIP2 in the lower leaflet, each corresponding to ∼2.5% of the bilayer composition. Pairwise nonbonded interactions between NAT and protein residues were analyzed for three replicas of 600 ns production runs (for the active ‘S4 up’ state) and four replicas of 1.2 μs production runs (for the resting ‘S4 down’ state) (Fig. 4A and Suppl. Fig. 5). Of all 262 residues, the outermost gating charge in S4, R227, clearly had the strongest electrostatic interaction with the anionic headgroup of NAT in all active state simulations. A secondary binding region was formed from K146 in the VSD and K284 in the pore domain (Fig. 4A-B). The accumulated sampling regions of NAT around KCNQ3 are visualized in Fig. 4C, with the highest NAT headgroup and tail density shown in 3D volume map. R227, K146, and K284 were also the strongest binding residues in the resting state simulations, but their interactions with NAT were weaker, even though the simulations were twice as long (Suppl. Fig. 5B).

**Figure 4.**
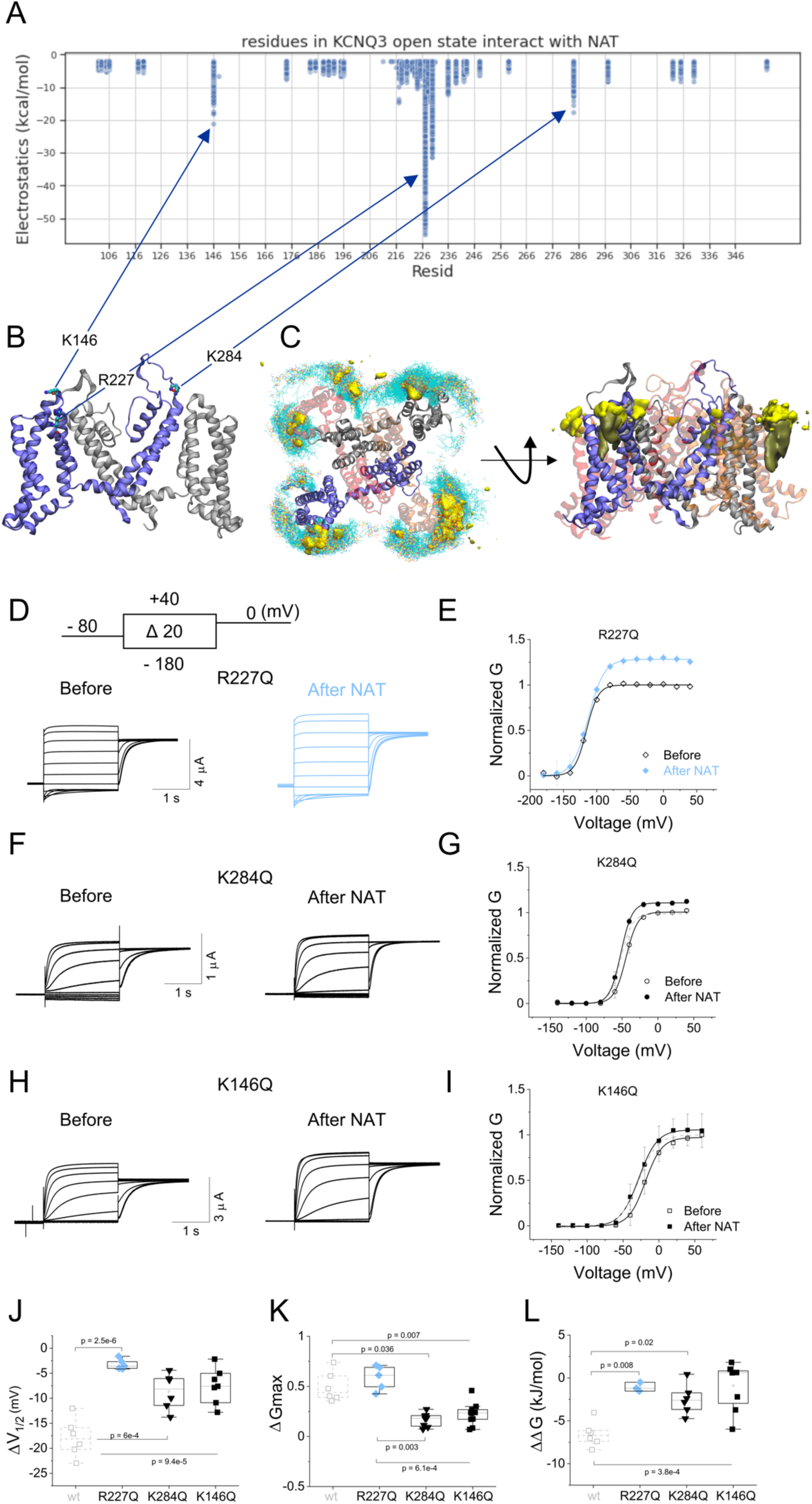
PUFAs primarily target the KCNQ3 voltage sensor. (*A*) Pairwise electrostatic interaction between KCNQ3 active (S4 up) state and NAT. Data points represent the electrostatic interaction from one of 200 snapshots over 2 μs. Only favorable interactions < –2 kcal/mol are shown for clarity. (*B*) Top three binding residues illustrated on one of the KCNQ3 subunits. (*C*) Volumetric map of NAT density (headgroup in yellow and tail in cyan). In the view from above (left), positions of NAT within 10 Å of KCNQ3 from 800 snapshots over 2 μs were overlapped. (*D, F, and H*) Representative current traces from KCNQ3-A315T-R227Q (D), KCNQ3-A315T-K284Q (F), and KCNQ3-A315T-K146Q (H) channels in the absence (before) or presence (after) of 25 μM NAT for the indicated voltage protocol. (*E, G, and I*) Steady-state G(V) relationships (solid lines from a Boltzmann fit) obtained from recordings in panels D, F, and H normalized to peak conductance before (open symbols) and after (closed symbols) NAT application, respectively. Dashed lines represent respective normalized G(V) relationships. (*J*-*L*) Summary data for ΔV_1/2_ (J); ΔGmax (K); and 11G (L) induced by 25 μM NAT for the indicated KCNQ3 mutation. Gray dashed boxes represent NAT–induced change of ‘wt’ KCNQ3-A315T in (J) ΔV_1/2_, (K) ΔGmax, and (L) 11G for comparison. Values are given in Table 2. Data represent mean ± SEM; n = 5–8. Statistical significance determined using one-way ANOVA and Bonferroni’s post hoc test to compare the NAT–induced change in ΔV_1/2_, ΔGmax, and 11G. p < 0.05; only significant differences are shown.

We tested the importance of R227 for NAT binding using the charge neutralizing mutation R227Q (Fig. 4B). In contrast to its effect on wt KCNQ3 channels, 25 μM NAT was unable to shift the KCNQ3-R227Q G(V) curve or change the energy required to open these channels (Fig. 4D, E, J, L, and Table 2). This suggests that S4, particularly R227, is important for mediating the effect of NAT on voltage dependence and therefore that this domain may form the primary PUFA binding site. However, because NAT remained able to increase Gmax in KCNQ3-R227Q channels (Fig. 4E, K, and Table 2), we wondered whether K146 and K284 may form a second binding site for this PUFA. We therefore made neutralizing mutations of K146 in the S1-S2 loop of the VSD and K284 at the top of S5 in the pore domain (Fig. 4B). Compared to wt, the K284Q and K146Q mutations significantly reduced the ability of NAT to left shift the G(V) curve and lower the energy required to open the channels (Fig. 4F-J, L, and Table 2). Furthermore, both K284Q and K146Q channels reduced the ability of NAT to increase Gmax by ∼45% and ∼50%, respectively (Fig. 4F-I, K and Table 2). These data indicate that K284 and K146 are important for mediating the effect of NAT on the voltage dependence and Gmax of KCNQ3 channels.

Previous work in the related cardiac I_KS_ (KCNQ1/KCNE1) channel showed that PUFAs increased Gmax via an electrostatic interaction between their negatively charged headgroup and a conserved positively charged residue in the S6 helix of KCNQ1 (K326)^46^. We therefore tested whether the homologous residue in KCNQ3, R330, is involved in NAT binding by mutating it to glutamine. Because homomeric R330Q mutant channels alone did not express, we co-injected wt KCNQ2 to partially rescue their functionality^15^ and create heteromeric KCNQ2/KCNQ3-R330Q channels (wtQ2/Q3-R330Q in a 1:1 ratio) (Suppl. Fig. 6). 25 μM NAT caused a 1.8-fold increase in Gmax in wt heteromeric KCNQ2/3 channels (Suppl. Fig. 6A-B and Suppl. Table 1). However, the effect of NAT on Gmax was reduced by ∼45% in heteromeric channels bearing the R330Q mutation (wtQ2/Q3-R330Q) (Suppl. Fig. 6C-D and Suppl. Table 1). R330Q did not completely abolish the ability of NAT to increase Gmax, likely because KCNQ2 also contains a conserved positively charged residue in its S6 helix (R291) that is homologous to R330 in KCNQ3. Heteromeric channels bearing a neutralizing mutation of R291 in KCNQ2 (Q2-R291Q/wtQ3) displayed a ∼27% reduction compared to wt KCNQ2/3 channels (Suppl. Fig. 6E-F and Suppl. Table 1). Moreover, heteromeric channels bearing both R291Q and R330Q mutations reduced the Gmax effect of NAT by ∼70%, but they did not completely abolish it (Suppl. Fig. 6G-H and Suppl. Table 1).

Taken together, these data suggest that NAT activates KCNQ3 channels involving two distinct regions of the channel. R227 on the N-terminal region of S4, and to a lesser extent K146 in S1 and K284 in S5, are involved in the shift in voltage dependence (leftward G(V) shift) mediated by NAT. In contrast, the extracellular regions of S1, S5, and S6 (K146, K284, and R330 respectively, and R291 in KCNQ2 in heteromeric KCNQ2/KCNQ3 channels), but not S4, are responsible for the increase in current amplitude (Gmax) induced by NAT.

### PUFAs rescue NDD-associated KCNQ3 mutations

Because NAA^+^ and NAT were the most potent inhibitor and activator, respectively, in our screen on KCNQ3 channels, we tested whether they could rescue the functionality of R227Q– and R236C-containing channels. Bath perfusion of 25 μM NAA^+^ modestly restored the G(V) curve of KCNQ3-R227Q channels to more positive voltages (Fig. 5A, B, E, and Table 2). NAA^+^ also reduced both Gmax (by ∼15%) and the amount of potassium current at negative voltages (Fig. 5A-B, F-G, and Table 2). Increasing NAA^+^ to 50 μM resulted in a larger reduction of Gmax (by ∼30%), but had no additional effect on the G(V) curve, in KCNQ3-R227Q channels (Fig. 5B and Table 2). Consistent with its inhibitory effect, NAA^+^ increased the energy required to open R227Q channels (Fig. 5H and Table 2), slowed the time course of current activation, and accelerated the time course of current deactivation (Suppl. Fig. 7A, B, E, and Table 2).

**Figure 5.**
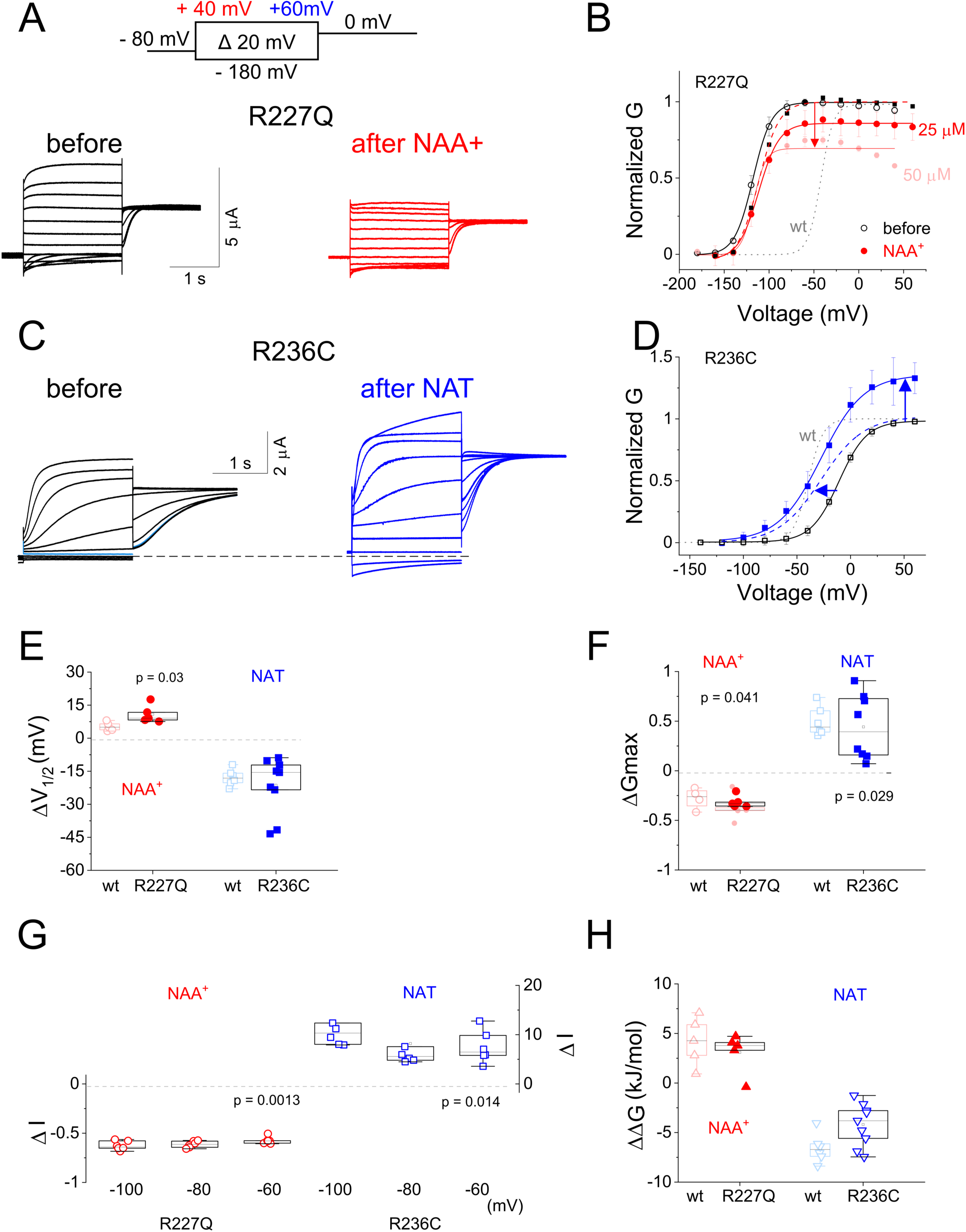
PUFAs rescue NDD-associated mutations. (*A*, *C*) Representative current traces from R227Q (A) and R236C (C) channels in the absence (before) or presence (after) of 25 μM NAA^+^ (A, red) and NAT (C, blue) for the indicated voltage protocol. (*B*, *D*) Steady-state G(V) relationships (solid lines from a Boltzmann fit) obtained from recordings in panels A and C normalized to peak conductance before (open symbols) and in the presence (closed symbols) of 25 μM and 50 μM NAA^+^ (B) and 25 μM NAT (D). Dotted gray lines represent G(V) curves for ‘wt’ KCNQ3-A315T channels and dashed lines represent respective normalized G(V) relationships after application of the indicated PUFA. (*E*-*H*) Summary data for ΔV_1/2_ (E); ΔGmax (F); 1I_–60_ (G); and 11G (H) induced by 25 μM NAA^+^ (red) in R227Q channels and 25 μM NAT (blue) in R236C channels. Dashed lines represent DMSO-induced changes for comparison. Data represent mean ± SEM, n = 5–9. Values are given in Table 2. Statistical significance determined using Student’s T-test to compare the PUFA–induced change in ΔV_1/2_, ΔGmax, 11G, 1I_–60_, and time to I_50%_ relative to mock application of a solution containing only vehicle (DMSO) for the indicated mutation. p < 0.05; only significant differences are shown.

In contrast, treatment of R236C-bearing channels with 25 μM NAT restored the G(V) curve to more negative voltages (Fig. 5C, D, E, and Table 2). NAT also increased Gmax by ∼30% and potentiated potassium currents by more than 7-fold at physiologically relevant voltages (Fig. 5F, G, and Table 2). Moreover, NAT decreased the energy required to open R236C channels by ∼4 kJ/mol (Fig. 5H and Table 2), accelerated the time course of current activation, and slowed the time course of current deactivation (Suppl. Fig. 7C-E, and Table 2). Together, these data indicate that NAA^+^ can partly rescue the GOF phenotype induced by R227Q and, importantly, that NAT can rescue the LOF phenotype induced by R236C so that this variant more closely resembles the properties of wt KCNQ3 channels.

### PUFAs act by modifying S4 movement and channel gating

To determine whether the ability of PUFAs to rescue the biophysical properties of R227Q and R236C variants was due to an effect on S4 movement, gate function, or coupling between S4 movement and gate opening (Fig. 6), we used VCF to measure S4 movement and channel gating simultaneously.

**Figure 6.**
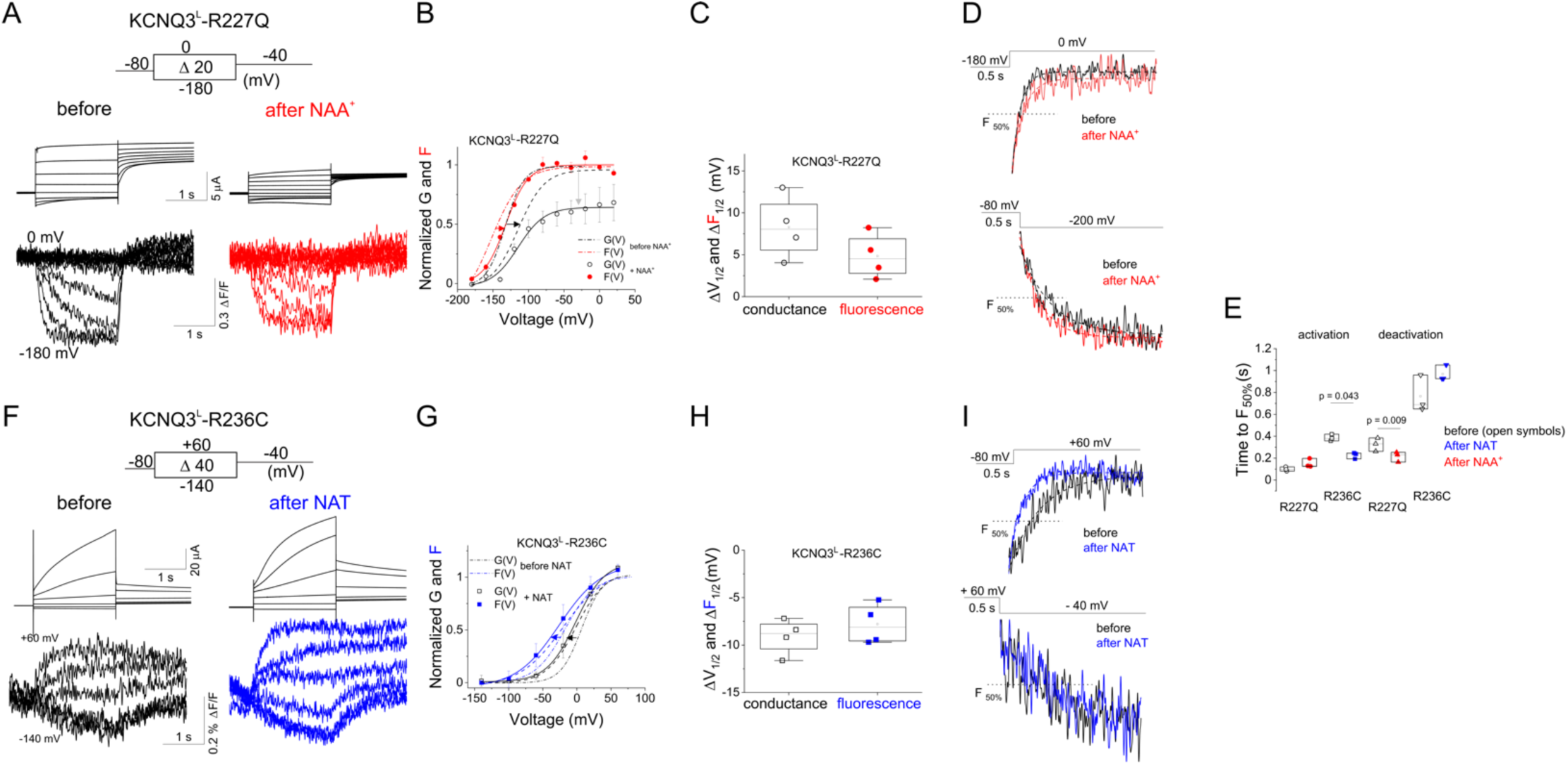
PUFAs act by modifying S4 movement and channel gating. (*A, F*) Representative current (top) and fluorescence (bottom) traces from Alexa-488–labeled KCNQ3^L^-R227Q (R227Q) (A) and KCNQ3^L^-R236C (R236C) (F) channels in the absence (before) or presence (after) of 25 μM NAA^+^ (red) and NAT (blue) for the indicated voltage protocol. (*B, G*) Steady-state G(V) relationships (black circles and black solid line from a Boltzmann fit) and F(V) relationships obtained from recordings in panels A (red circles and red solid line from a Boltzmann fit) and F (blue squares and blue solid line from a Boltzmann fit), normalized to peak conductance and fluorescence before PUFA application (open symbols). Dotted and dashed-dotted lines represent G(V) and F(V) curves for untreated R227Q (B) and R236C (G) channels, respectively. Dashed lines represent respective normalized G(V) relationships after application of the indicated PUFA. (*C*, *H*) Summary data for ΔV_1/2_ (black) and ΔF_1/2_ (red and blue) obtained from recordings in panels B and G. (*D*, *I*) Representative time courses of fluorescence activation (top) and deactivation (bottom) in the absence (black) and presence of NAA^+^ (red) or NAT (blue) in KCNQ3^L^-R227Q (R227Q) (D) and KCNQ3^L^-R236C (R236C) (I) channels in response to the indicated voltage protocol. Dashed lines represent 50% of maximum fluorescence at the end of the pulse. (*E*) Time courses of fluorescence activation and deactivation in the absence (open black symbols) and presence (closed symbols) of NAA^+^ (red) or NAT (blue) are quantified as time to reach half maximum fluorescence at the end of the pulse (dashed lines). Data represent mean ± SEM, n = 3–5. Statistical significance determined using pair-sample Student’s t-test. p < 0.05; only significant differences are shown.

Similar to its effect on unlabeled R227Q, 25 μM NAA^+^ reduced the current carried by Alexa-488-labeled KCNQ3^L^-R227Q channels (Fig. 6A-B and Table 2) and shifted G(V) and F(V) curves to more positive potentials (Fig. 6B, C, and Table 2). NAA^+^ also moderately slowed the time course of current activation and did not alter the time course of current deactivation (Suppl. Fig. 8A-B and Table 2).

Furthermore, Alexa-488 fluorescence revealed that NAA^+^ did not alter the time course of fluorescence signal (S4) activation, but accelerated the time course of fluorescence signal deactivation, compared to untreated KCNQ3^L^-R227Q channels (Fig. 6D, E, and Table 2). Analysis of Gibbs free energy showed that NAA^+^ increased the energy required to open labeled KCNQ3^L^-R227Q channels, but moderately increased the energy required for S4 movement (Table 2) and slightly decreased the amplitude of Alexa-488 fluorescence in labeled KCNQ3^L^-R227Q channels (Fig. 6A). The mechanism by which NAA^+^ quenches fluorescence is unclear, but it is plausible that it induces a rearrangement of the channel that positions a nearby quenching residue, like tryptophan, closer to the fluorophore.

In contrast, treatment with NAT led to a leftward shift of both the G(V) and F(V) relationships (Fig. 6F-H and Table 2), and accelerated the time course of current activation (without affecting deactivation), in KCNQ3^L^-R236C channels (Suppl. Fig. 8A and Table 2). Moreover, NAT accelerated the time course of fluorescence activation (S4 movement), but did not affect the time course of fluorescence deactivation, in KCNQ3^L^-R236C channels (Fig. 6I, E, and Table 2). Consistent with its positive effect on activity, NAT decreased the energy required to move S4 in KCNQ3^L^-R236C channels (Table 2).

Taken together, our VCF data suggest that NAT is able to effectively activate KCNQ3-R236C channels by altering S4 movement and channel gating. NAA^+^ had a more moderate effect on KCNQ3-R227Q channels, likely because the mutation itself removed the positive charge at position 227 that is important for NAA^+^ to shift the G(V) curve.

## Discussion

Pathogenic variants in KCNQ3 (M-channel) subunits are associated with NDDs, including devastating developmental and epileptic-encephalopathy^10, 11^. However, the impact these KCNQ3-channelopathies have on M-channel function and NDD pathogenesis remain poorly defined and existing treatments are often ineffective or have serious side effects limiting their utility^26, 47^. We have investigated the functional consequences of two mutations in the S4 helix of KCNQ3 channels that are linked to NDDs and autism. We found that R227Q, a neutralization of the first gating charge, cause gain-of-function by impairing S4 movement without affecting the gate. In contrast, R236C, a neutralization of the third gating charge, cause loss-of-function by altering S4-to-gate coupling.

Furthermore, we showed that two PUFAs, NAA^+^ and NAT^-^, can rescue the wt function of R227Q and R236C phenotypes, respectively. The mechanisms by which PUFAs act on KCNQ3 channels and KCNQ3-bearing pathogenic variants were examined using all-atom MD simulations and electrophysiological analyses. Our data suggest that the outermost positive gating charge in the voltage sensor (R227) and positively charged residues in the extracellular regions of S1, S5, and S6 (K146, K284, and R330), but not the retigabine– and ICA-binding pockets, are critical for mediating PUFAs effect on voltage-dependent shift and Gmax. The R227Q mutant (and to a lesser extent K284Q and K146Q) abolished the effect of NAT on voltage dependence, supporting the hypothesis that an electrostatic interaction between the negative charge of the taurine head group of PUFA and the positive charge of arginines in S4 of KCNQ3. Our data provide a proof of principle that PUFAs can at least partially restore wt functionality in KCNQ3 channels bearing NDD-associated variants, laying the groundwork for future development of more potent molecules to correct NDD-associate alterations in KCNQ function.

The molecular basis of NDDs is largely unknown. Recent studies have provided evidence linking voltage-gated ion channels, including KCNQ channels, with the pathophysiological features of NDDs^48, 49^. The KCNQ3-R227Q mutation has been reported to cause sleep-activated near-continuous multifocal spikes^9^ as well as autism without neonatal seizures^10, 16^. Compared to wt homomeric KCNQ3 channels, homomeric channels carrying the R227Q mutant exhibit a GOF phenotype that includes enhanced current density and a prominent leftward shift in the G(V) curve. These effects were partially attenuated upon co-expression of wt KCNQ2 and KCNQ3, in agreement with previous observations^9^. Our VCF data showed that the R227Q mutation shifts the F(V) curve (representing S4 movement) to similar voltages as the G(V) curve (representing gate movement). In addition, the time course of S4 movement and channel gating in R227Q mutant channels are closely correlated, as they are in wt KCNQ3 channels. These results indicate that mutating the first gating charge (R227Q) of KCNQ3 alters channel function by disturbing S4 activation and not by affecting the relationship between S4 movement and channel gating.

We recently showed that neutralization of the first gating charge in the related KCNQ2 channel (R198Q) resulted in a similar leftward shift of the G(V) curve and an overlap of this curve with the F(V) relationship^50^. This close correlation between voltage dependence of pore opening and S4 movement was reflected in the similar time courses of ionic currents and fluorescence signals^50^, as we observed for KCNQ3-R227Q in the current study. Previous studies using systematic mutagenesis and molecular modeling of S4 in other members of the KCNQ family demonstrated that disease-associated neutralizing mutations of the first and the second gating charge of S4 caused a leftward shift in the voltage G(V) curve^51, 52, 53^, similar to that observed in KCNQ3-R227Q channels. These studies proposed that the probable mechanism underlying gain of voltage sensitivity in these disease-causing mutations was likely due to destabilization of the resting state configuration of S4, which would explain other GoF effects. Furthermore, our earlier work showed that neutralization of the second arginine within S4, which has previously been linked to NDD^9^, disrupted KCNQ3 channel function by shifting the voltage dependence of S4 movement to extreme negative potentials and driving the open/closed transition of the gate to very negative potentials^42^. Therefore, it appears that basic residues in the N-terminal part of S4 (first and second gating charges) are crucial for stabilizing the resting state of the voltage sensor and keeping the channel closed; a mechanism that seems to be conserved among KCNQ channels^50, 53, 54, 55^.

In contrast to mutations in the N-terminal part of S4, mutations of charged residues in the C-terminal region of S4 (third, fourth and fifth gating charges) have been reported to reduce KCNQ channel activity (LOF), primarily by shifting the G(V) curve toward more depolarized voltages and either slowing the kinetics of activation or accelerating the kinetics of deactivation^16, 56^. Motivated by these findings, we used VCF to examine how the R236C mutation in KCNQ3 affected channel activity. In contrast to the R227Q, the R236C variant caused a dissociation of S4 movement and channel opening, which we observed as a more hyperpolarized F(V) curve than G(V) curve. Thus, although both S4 movement and gate opening were affected by the R236C mutation, S4 movement changed less than channel opening, suggesting that R236C altered S4-to-gate coupling. In a previous study^50^, we observed a similar result for the epilepsy-associated R214W mutation in homomeric KCNQ2 channels, suggesting similarities between KCNQ2 and KCNQ3 gating mechanisms. However, KCNQ2-R214 is in the loop connecting S4 and the S4-S5 linker and is thought to coordinate the binding of PIP2 – a phospholipid that regulates channel gating^50^ – whereas KCNQ3-R236 is in the S4 segment – two helical turns above the homologous R214 residue – and is therefore unlikely to form part of the PIP2 binding pocket.

Our study also identified PUFAs that modulate KCNQ3 channel activity restoring functionality in NDD-associated KCNQ3 channel variants. The two most effective PUFAs –NAT^−^ and NAA^+^– modified voltage sensitivity, maximum current amplitude, and both activation and deactivation kinetics. Previous studies in shaker^57, 58^ and cardiac KCNQ1^59^ channels showed that charged PUFA analogs form electrostatic interactions with positively charged arginines in the N-terminal region of S4 to activate these channels. Our finding that the R227Q mutant reduced KCNQ3 channel sensitivity to NAT^−^ and NAA^+^ suggests that the negatively charged taurine group of NAT^−^ and the positively charged amino group of NAA^+^ may attract and repel the guanidinium group of R227 to stabilize and destabilize the activated state of S4, respectively. This would explain why NAT^−^ functions as a channel activator and NAA^+^ as an inhibitor, and furthermore, why NAA^+^ had a minimal effect on S4 movement and voltage dependence but more effectively accelerated current deactivation and reduced maximum current amplitude. NAA^+^ caused only a small change in the time course of S4 movement, likely due to the absence of electrostatic repulsion between R227Q (in the S4) and NAA^+^.

VCF showed that NAT^−^ shifted the properties of R236C mutant channels towards those of wt KCNQ3 channels, including the voltage dependence and time course of S4 movement and channel opening. Since R236C changes the coupling of voltage sensor movement and gate opening, it is tempting to speculate that NAT^−^ may impact both the S4 and the gate, as previously suggested for KCNQ1^60^. In KCNQ3, we found that NAT^−^ interacts with R227 (N-terminal region of S4), a position distinct from the retigabine and ICA binding sites, resembling the mechanism of action proposed for endocannabinoids in KCNQ2/3 channels^29^. It is possible that the negative taurine moiety of NAT^−^ may be attracted to the charged residues of S4, particularly R227, thereby stabilizing the activated state of S4 via electrostatic mechanisms. This would indirectly exert a restoration of the voltage dependence of KCNQ3 channel opening and restore function of the KCNQ3-R236C mutant.

In addition of NAT’s interaction with S4 of KCNQ3, MD simulations also revealed a secondary interaction with K146 and K284, located in the extracellular loops of S1 and S5, between the VSD, lipids, and extracellular side of the pore. These data are consistent with our findings that K146Q and K284Q reduce the effect of NAT^−^ on Gmax and voltage dependence, indicating that NAT^−^ could also form electrostatic interactions with positively charged residues on the pore domain. Previous MD simulations of PUFAs have identified electrostatic interactions with the pore and VSD domains of KCNQ1^61^. Interestingly, our MD simulations show that NAT^−^ binds more strongly to S4 in the activated (up) state than the resting (down) state, likely due to a greater exposure of the R227 binding site to the membrane surface in the up state. Such state-dependent binding suggests that NAT^−^ stabilizes the activated conformation of S4, consistent with the slower deactivation kinetics of KCNQ3 in the presence of NAT^−^.

Our study reveals that the molecular mechanisms underlying KCNQ channelopathies could be heterogeneous, and that predicting their functional consequences could be challenging. For instance, it is often assumed that mutations affecting VSDs change the voltage-dependence of opening and result in a concomitant alteration of opening probability, whereas mutations in the pore region hamper permeability. However, our data shows that, although both R227 and R236 are located in S4 and are linked to similar NDD phenotypes^11^, the mechanisms by which mutations of these residues perturb channel gating are different. An understanding how mutations affect different regions of a channel is critical for the design of appropriate therapeutic strategies to treat these disabling conditions while minimizing potential side effects.

Most NDD-associated missense variants in the KCNQ3 channel alter arginine residues in the VSD^9, 11^. Although the majority of these alterations that have been characterized lead to GOF^9^, likely by direct disturbance of S4, some variants (including R236C) lead to LOF due to dissociation of S4 movement and gate opening. The antagonistic biophysical properties of NAA^+^ and NAT can compensate for the enhanced and reduced activity of NDD variant-bearing channels. The divergent molecular properties of these arachidonic acid derivatives could therefore be used as scaffold molecules to design more potent, and patient specific, drugs to treat neurological disorders associated with KCNQ channel defects.

## Methods

### Chemicals

(μ-3) 4,7,10,13,16,19-all-cis-docosahexaenoic acid (DHA); (μ-6) 9, 12-cis-linoleic acid (LA); (μ-6) 5,8,11,14-all-cis-arachidonic acid (AA); 4,7,10,13-cis-arachidonoyl ethanolamine (O-AEA); 9, 12-cis-linoleoyl-glycine (Lin-gly); and 5,8,11,14-all-cis-N-arachidonoyl taurine (NAT) were purchased from Cayman Chemicals Inc (Ann Arbor, MI, USA). 5,8,11,14-cis-N-arachidonoyl amine (NAA^+^) was synthesized at Linkoping University, Sweden. Alexa Fluor 488 C5-maleimide was purchased from Thermo Fisher Scientific (Waltham, MA, USA). All other chemicals were obtained from Sigma-Aldrich (St. Louis, MO, USA). PUFAs were kept at −20°C as 50 mM stock solutions in DMSO (NAT) or ethanol (NAA^+^) and diluted shortly before experiments as previously described^57^. Control and test solutions were added to the recording chamber using the Rainin Dynamax peristaltic pump-driven (model RP-1) perfusion system.

### Molecular Biology

The full length human KCNQ3 construct (NCBI Reference Sequence: NP_004510.1; GI:4758630) was synthesized (GenScript USA, Piscataway, NJ) and ligated between the BamHI and XbaI sites in the multiple cloning sites of into the pGEM-HE vector. This vector had been previously modified to contain a T7 promoter and 3’ and 5’ untranslated regions from the Xenopus β-globin gene ^42^. The Kozak consensus sequence (GCCACC) and the AgeI restriction site (ACCGGT) were added before the start codon (ATG) of the KCNQ3 gene. Point mutations were made in the KCNQ3 gene using the Quikchange XL site-directed Mutagenesis kit (Agilent) according to the manufacturer’s protocol. The correct incorporation of the specific variant was assessed by Sanger sequencing (sequencing by Genewiz LLC, South Plainfield, NJ). The RNA was synthesized in vitro using the mMessage mMachine T7 RNA Transcription Kit (ThermoFisher Scientific) from the linearized cDNA. mRNA (40-50nL) was injected into defolliculated *Xenopus leavis* oocytes (purchased from Ecocyte) using a Nanoject II nanoinjector (Drummond Scientific) and electrophysiological experiments were performed 2 to 5 days after injection.

### TEVC recordings

Two-electrode voltage-clamp (TEVC) recordings were performed as previously described^42, 62^. Regular ND96 solution for TEVC contained 96 mM NaCl, 2 mM KCl, 1 mM MgCl2, 1.8 mM CaCl2 and 5 mM HEPES (pH = 7.5). Ionic currents were recorded in TEVC using an OC-725C oocyte clamp (Warner Instruments), low-pass filtered at 1 kHz and sampled at 5 kHz. Microelectrodes were pulled using borosilicate glass to resistances from 0.3 to 0.5 MΩ when filled with 3 M KCl. Voltage clamp data were digitized at 5 kHz (Axon Digidata 1440A; Molecular Devices), collected using pClamp 10 (Axon Instruments).

### Voltage clamp fluorometry (VCF)

VCF experiments were carried out as previously reported^42^. Briefly, aliquots of 50 ng of mRNA coding for KCNQ3 or the KCNQ3 variant RNA were injected into *Xenopus laevis* oocytes. At 2-5 days after injection, oocytes were labeled for 30 min with 100 μM Alexa-488 maleimide (Thermo Fisher Scientific) in high [K^+^] solution (98 mM KCl, 1.8 mM CaCl_2_, 1 mM MgCl_2_, 5 mM HEPES, pH 7.05) at 4 °C, in the dark. The labelled oocytes were then rinsed three-to-five times in dye free ND96 and kept on ice before each recording to prevent internalization of labeled channels. Oocytes were placed into a recording chamber animal pole “up” in ND96 solution (pH 7.5 with NaOH) and electrical measurements were carried out in TEVC using an Axoclamp 900A amplifier (Molecular Devices). Microelectrodes were pulled to resistances from 0.3 to 0.5 MΩ when filled with 3 M KCl. Voltage clamp data were digitized at 5 kHz (Axon Digidata 1550B via a digital Axoclamp 900A commander, Molecular Devices) and collected using pClamp 10 (Axon Instruments). Fluorescence recordings were performed using an Olympus BX51WI upright microscope. Light was focused on the top of the oocyte through a 20× water immersion objective, 1 NA, after being passed through an Oregon green filter cube (41026; Chroma). Fluorescence signals were focused on a photodiode and amplified with an Axopatch 200B patch clamp amplifier (Axon Instruments). Fluorescence signals were low-pass Bessel-filtered (Frequency Devices) at 100–200 Hz, digitized at 1 kHz, and recorded using pClamp 10.

### MD simulations

The homology models of KCNQ3 channel transmembrane domain with S4 in the resting (down) state and active (up) state were created using the Swiss-model program (https://swissmodel.expasy.org/) based on the model of KCNQ1 channel in the resting state^63^ and KCNQ2 channel in its activated (S4 up) state and the pore in the closed state ^64^ (PDB: 7CR0), respectively. Each was embedded in POPC (palmitoyl oleoyl-phosphatidylcholine) (∼470 lipids per leaflet), ∼2.5% phosphatidyl-4,5-bisphosphate (PIP2) (23 in the lower leaflet), and N-Arachidonoyl taurine (NAT) (20 in the upper leaflet) using the membrane builder tool on CHARMM-GUI membrane builder^65^. All NAT lipids were placed randomly beyond 10 Å from KCNQ3. A TIP3P water layer of 30 Å thickness on the upper bilayer and lower leaflet containing 150mM of KCl was added in this system. CHARMM36 force field was used for protein^66, 67^, POPC and PIP2 lipids^68^, KCl, and TIP3P water^69^.

CHARMM general force fields (CGenFF)^70^ were used to prepare NAT parameters. NAMD2.13^71^ was used for equilibrium and production simulations. Cutoff for calculating van der Waals interactions and short-range electrostatic interactions was set at 12 Å and force-switched at 10 Å. Long-range electrostatic interactions were calculated using the particle mesh Ewald algorithm^72^.

### PUFA-KCNQ3 interaction analysis

Pairwise per-residue based Generalized Born (GB) energy decomposition was performed using idecomp=3 (adding 1-4 interactions to internal energy), igb=5, saltcon=0.150 options in MMPBSA.py (https://pubs.acs.org/doi/10.1021/ct300418h). The total pairwise energy is the sum of the van der Waals term, electrostatic term, generalized Born (GB) Polar solvation term, and Non-polar solvation term. 6 ns per snapshots were extracted from each trajectory for computing the interaction between each NAT lipid and one of 1048 protein residues. Three replicas of 600 ns runs were analyzed for the activated state. The resting state simulations were run with double length due to the weaker binding interactions. The sampling was determined to be sufficient when all replicates showed consistent ranking of the top three binding residues.

### Electrophysiology data Analysis

Data were analyzed with Clampfit 10 (Axon Instruments, Inc., Sunnyvale, CA), OriginPro 2021b (OriginLabs Northampton, MA), and Corel-DRAW Graphics Suite 2021 software. To determine the ionic conductance established by a given test voltage, a test voltage pulse was followed by a step to the fixed voltage of –40 mV (tail), and current was recorded following the step. To estimate the conductance g(V) activated at the end of the test pulse to voltage V, the current flowing after the hook was exponentially extrapolated to the time of the step and divided by the offset between –40 mV and the reversal potential. The conductance g(V) associated with different test voltages V in a given experiment was fitted by the relation:

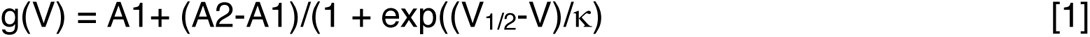

where A1 and A2 are conductances that would be approached at extreme negative or positive voltages, respectively, V_1/2_ the voltage that activates the conductance (A1+A2)/2, and κ is the slope factor in mV. Due to the generally different numbers of expressed channels in different oocytes, we compare normalized conductance, G(V):

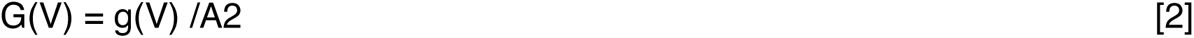

Fluorescence signals were corrected for bleaching and time-averaged over 10-40 ms intervals for analysis. The voltage dependence of fluorescence f(V) was analyzed and normalized (F(V)) using relations analogous to those for conductance (equations. 1 and 2).

To measure the effect of PUFAS, we stepped every 30 s from −80 mV to 0 mV for 5 s before stepping to −40 mV and back to −80 mV to ensure that the PUFA effects on the current at 0 mV reached steady state. We then used a voltage-step protocol to measure the current vs. voltage (I-V) relationship before and after the PUFA effects had reached steady state. Cells were held at −80 mV followed by a step from −140 (or as indicated in each figure) to + 40 mV (in 20 mV steps) followed by a subsequent voltage step to −40 mV (or 0mV) to measure tail currents before returning to the −80 mV holding potential.

The delta maximum conductance (βGmax) (or delta current at – 60 mV (△I_-60_)) was calculated by taking the difference between the Gmax in PUFAs (Gmax_PUFA_) (or I_-60 PUFA_) and Gmax without PUFAs (Gmax_0_) (or I_-60 0_), divided by Gmax_0_ (or I_-60 0_), as:

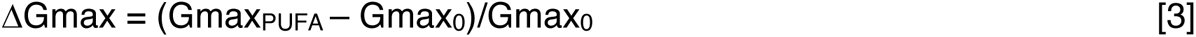

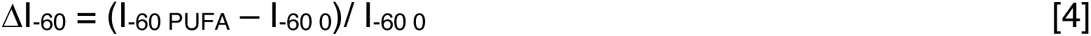

To estimate the effect of mutations (mut) relative to the wild type (wt) and the effect of PUFAs on Gibbs free energy (ΔΔG), the following relations were used:

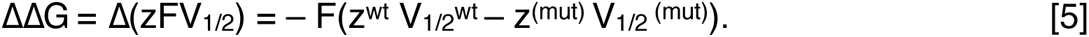

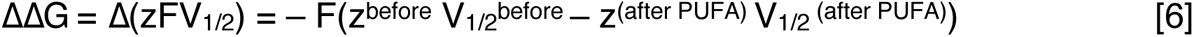

where F is Faraday’s constant, z (the slope; z = 25/s), and V_1/2_ (half-activation-voltage) were obtained from the mutation (mut), wild type (wt), control (before PUFA), and treated (after PUFA) G(V) curves^39, 40^. These analysis assumes a two-state model and tends to underestimate the z^73^. The calculated ΔΔG should therefore be seen as an approximation. To analyze the effect of PUFAs on current and fluorescence kinetics we used T50%, which for opening (and closing) it was defined as the time it takes to reach 50% of the maximum (and minimum) current (or fluorescence) level at the end of the indicated depolarizing (opening or activated S4) and hyperpolarizing (closed or resting S4) pulse.

### Statistics

All experiments were repeated 4 or more times from at least three batches of oocytes. Pairwise comparisons were achieved using ANOVA and Bonferroni’s post hoc test or Student’s t test as indicated in each figure. Data are represented as mean ± s.e.m (standard error of mean) and “n” represents the number of experiments.

## Acknowledgements

We thank Dr. Peter Larsson, for helpful comments on the manuscript. This work was supported by the National Institutes of Health (1R01NS110847) to RB-S.

---

**Supplementary Figure 1.**
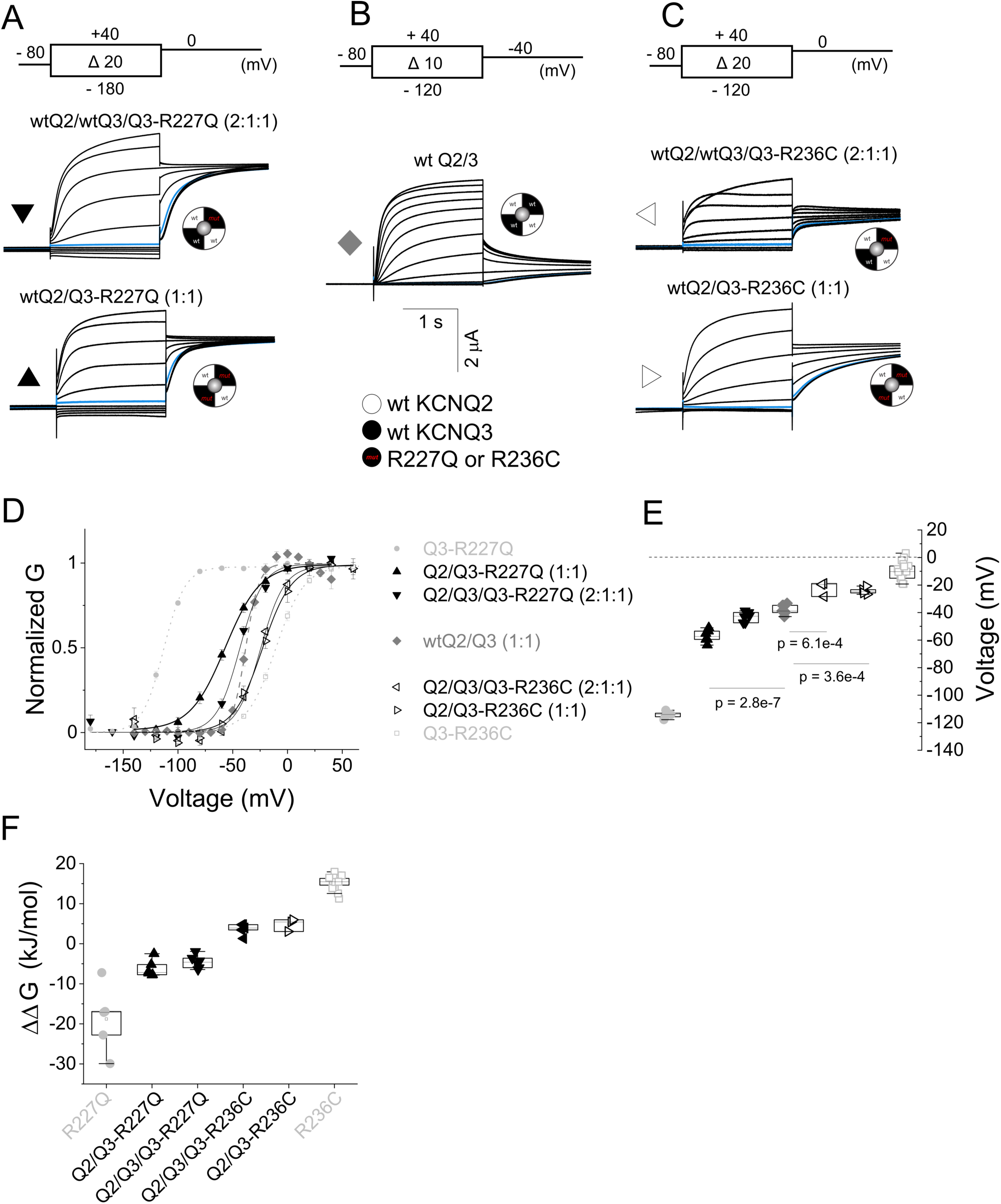
Biophysical phenotype of heterozygous NDD R227Q and R236C mutants. (*A, C*) Representative current traces from channels assembled from KCNQ2, KCNQ3, and KCNQ3-R227Q (A) or KCNQ3-R236C (C) expressed in a (top) 2:1:1 ratio (wtQ2/wtQ3/Q3-R227Q or wtQ2/wtQ3/Q3-R236C) and in a (bottom) 1:1 ratio (wtQ2/Q3-R227Q or wtQ2/Q3-R236C) to mimic heteromeric channels in a heterozygous patient case. (*B*) Representative current traces from wild type heteromeric KCNQ2/3 (wt Q2/3) channels expressed at a 1:1 ratio. (D) Extrapolated tail conductance from panels A-C were normalized and plotted against test voltages to create G(V) curves. Lines represent the fitted theoretical voltage dependencies (eqs. 1 and 2). (*E-F*) Summary data for (E) G(V) midpoints using Boltzmann fits from panel (D) and (*F*) Gibb free energy (11G). Midpoints of voltage activation and 11G values for each channel are shown in Suppl. Table 1. Data presents the mean ± SEM. Statistical significance was determined using ANOVA and Bonferroni’s post hoc test, and significance level was set at p < 0.05.

**Supplementary Figure 2.**
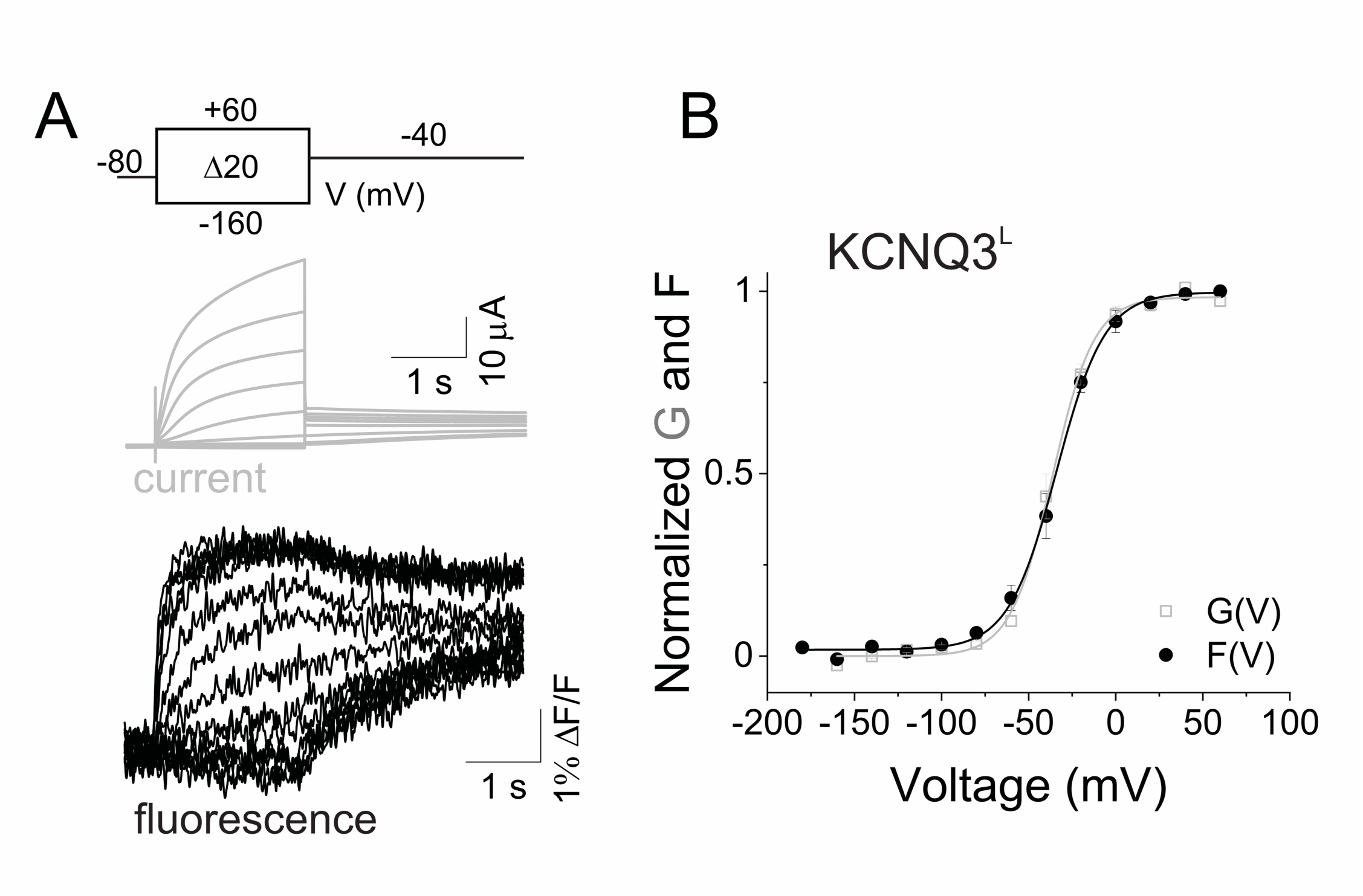
**Labeled KCNQ3-A315T-Q218C (KCNQ3^L^) channels track S4 movement**. (*A*) Representative current (gray) and fluorescence (black) traces from Alexa-488– labeled KCNQ3-A315T-Q218C (KCNQ3^L^) channels for the indicated voltage protocol. (*B*) Normalized G(V) (squares and solid gray line from a Boltzmann fit) and F(V) (circles and black line from a Boltzmann fit) curves from KCNQ3^L^ channels. The midpoints of activation of the fits are shown in Table 1. Data represents the mean ± SEM; n = 13.

**Supplementary Figure 3.**
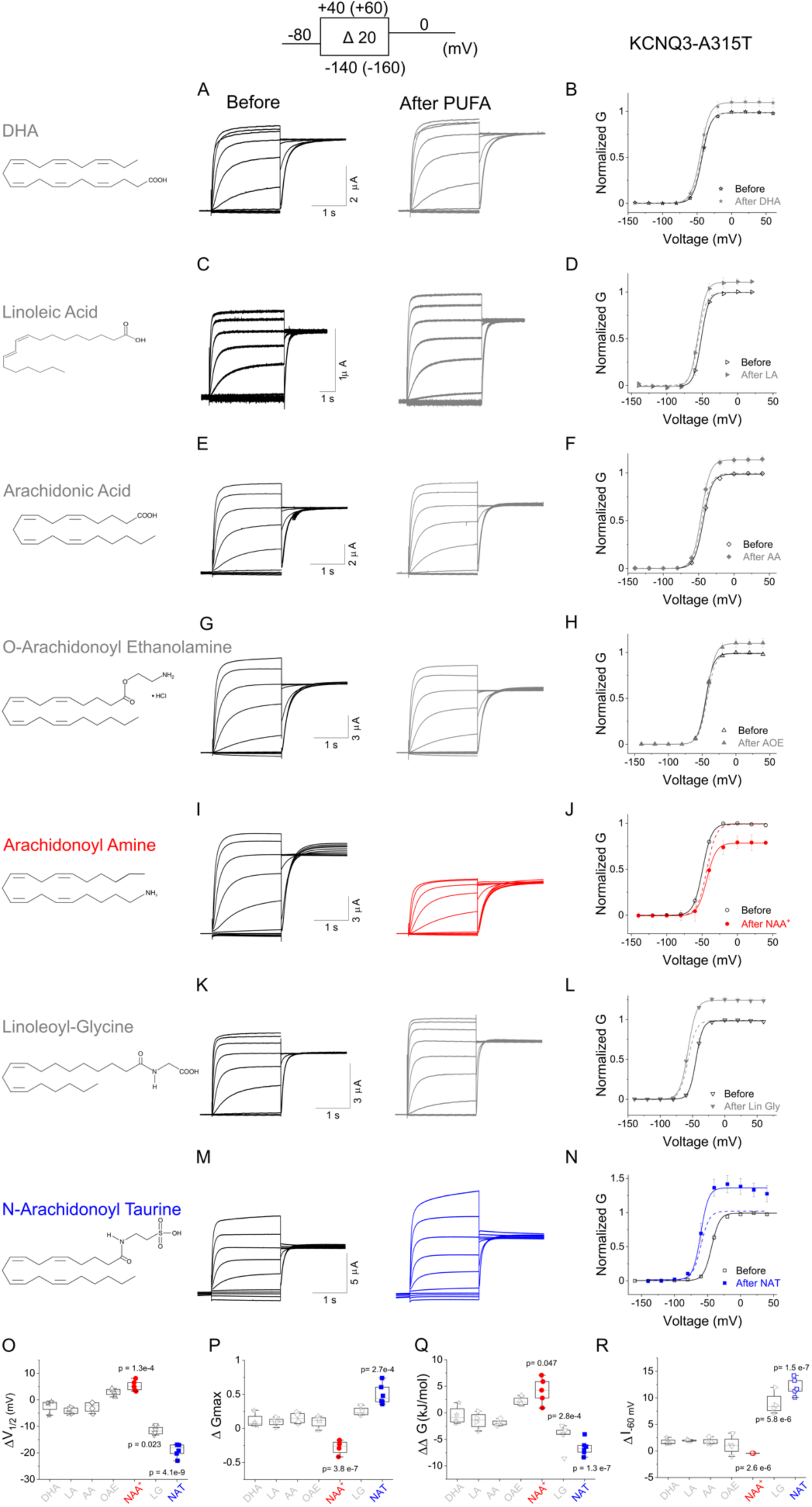
Effect of PUFA and PUFA analogs on KCNQ3 channel gating. (*A, C, E, G, I, K, and M*) Representative current traces from KCNQ3-A315T channels in the absence (before) or presence (after) of 25 μM of the indicated PUFAs for the indicated voltage protocol. Molecular structures of each PUFA: docosahexaenoic acid (DHA), linoleic acid (LA), arachidonic acid (AA), arachidonoyl ethanolamine (O-AEA), N-arachidonoyl amine (NAA^+^), linoleoyl-glycine (Lin-gly), and N-arachidonoyl taurine (NAT) are shown on the left. (*B, D, F, H, J, L, and N*) Steady-state conductance/voltage relationships, G(V)s, (solid lines from a Boltzmann fit) obtained from recordings in panels *A, C, E, G, I, K, and M* normalized to peak conductance before PUFA application (open symbols). G(V) relationships (solid lines from a Boltzmann fit) in the presence of PUFAs shown as closed symbols and dashed lines represent their respective G(V) relationships. (O-R) Summary data for shifts in half-activation voltage (ΔV_1/2_) for each G(V) (O), increase in Gmax (ΔGmax) (P), 11G (Q), and relative change in potassium current at –60 mV (1I_–60_) (R) induced by 25 μM PUFAs on KCNQ3 channels. Values for V_1/2,_ 1V_1/2_, 1Gmax (eq 3), 11G (eq 6), 1I_-60_ (eq 4), and time to I_50%_ given in Suppl. Table 2. Data represents the mean ± SEM; n = 6–11. Statistical significance was determined using one-way ANOVA and Bonferroni’s post hoc test to compare the PUFAs–induced changes in half-activation V_1/2_ (ΔV_1/2_), ΔGmax, 11G, and 1I_–60_ relative to mock application of a solution containing only vehicle (DMSO). Significance level was set at p < 0.05 and only significant differences are shown.

**Supplementary Figure 4.**
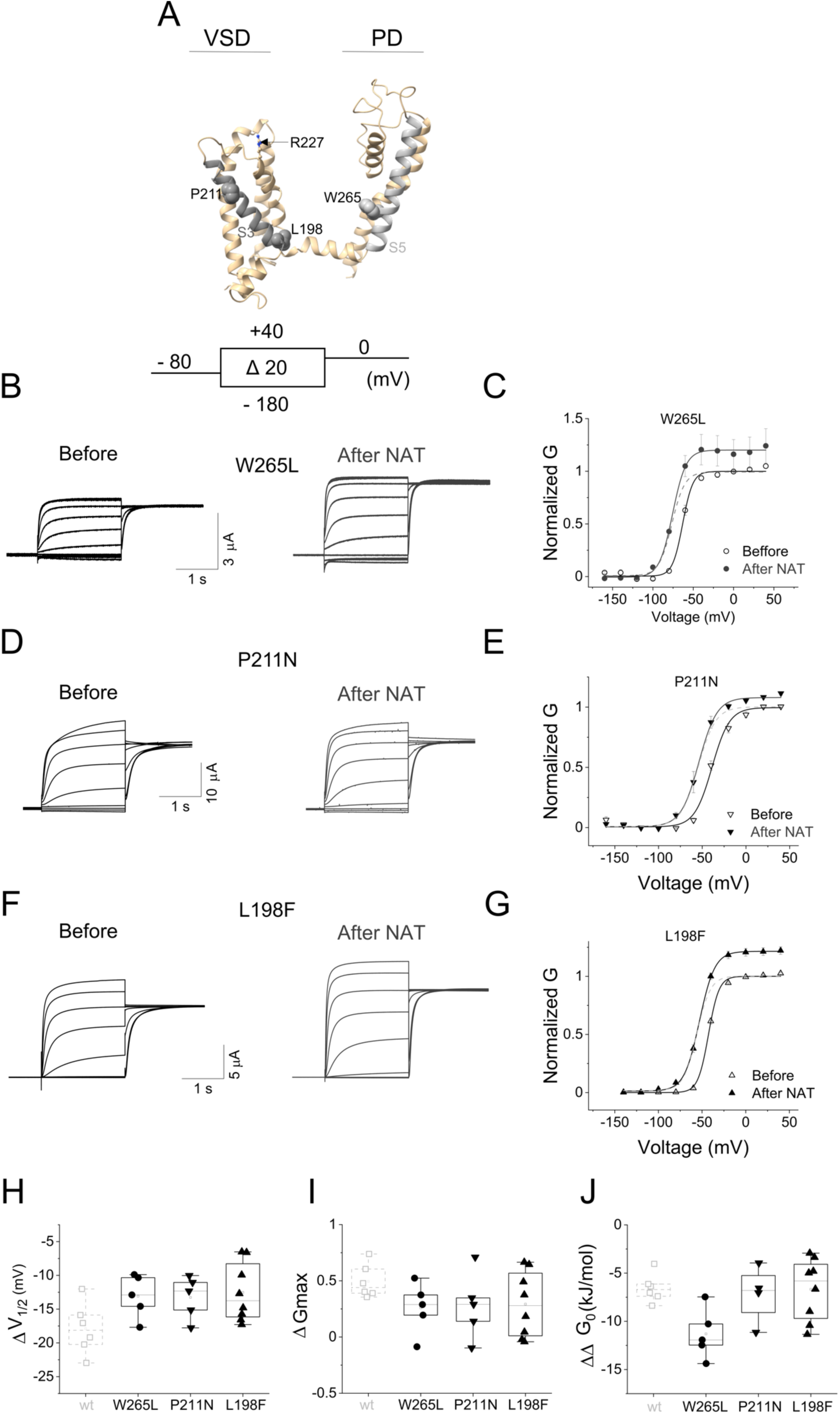
NAT and KCNQ channel openers have distinct binding sites. (*A)* KCNQ3 homology model of the activated (S4 up) and closed (S4 down) state using the Swiss-model program (https://swissmodel.expasy.org/) based on the cryo-EM structure of KCNQ2 (PDB code: 7CR0). One subunit is shown as ribbons with key amino acid residues as sticks. VSD, voltage sensing domain; PD, pore domain. Image generated using UCSF ChimeraX, version 1.1 (2020-10-07). (*B, D, and F*) Representative current traces from KCNQ3-A315T-W265L (B), KCNQ3-A315T-P211N (D), and KCNQ3-A315T-L198F (F) channels in the absence (before) or presence (after) of 25 μM NAT for the indicated voltage protocol. (*C, E, and G*) Steady-state G(V) relationships (solid lines from a Boltzmann fit) obtained from recordings in panels *B, D, and F* normalized to peak conductance. Dashed lines represent their respective normalized G(V) relationships. (*H*-*J*) Summary data for ΔV_1/2_ (H); ΔGmax (I); and 11G (J) induced by 25 μM of NAT for the indicated KCNQ3 mutation. Data represent mean ± SEM; n = 5–8. Values are given in Suppl. Table 2. Statistical significance determined using one-way ANOVA and Bonferroni’s post hoc test to compare the NAT– induced change in ΔV_1/2_ and 11Go. p < 0.05.

**Supplementary Figure 5.**
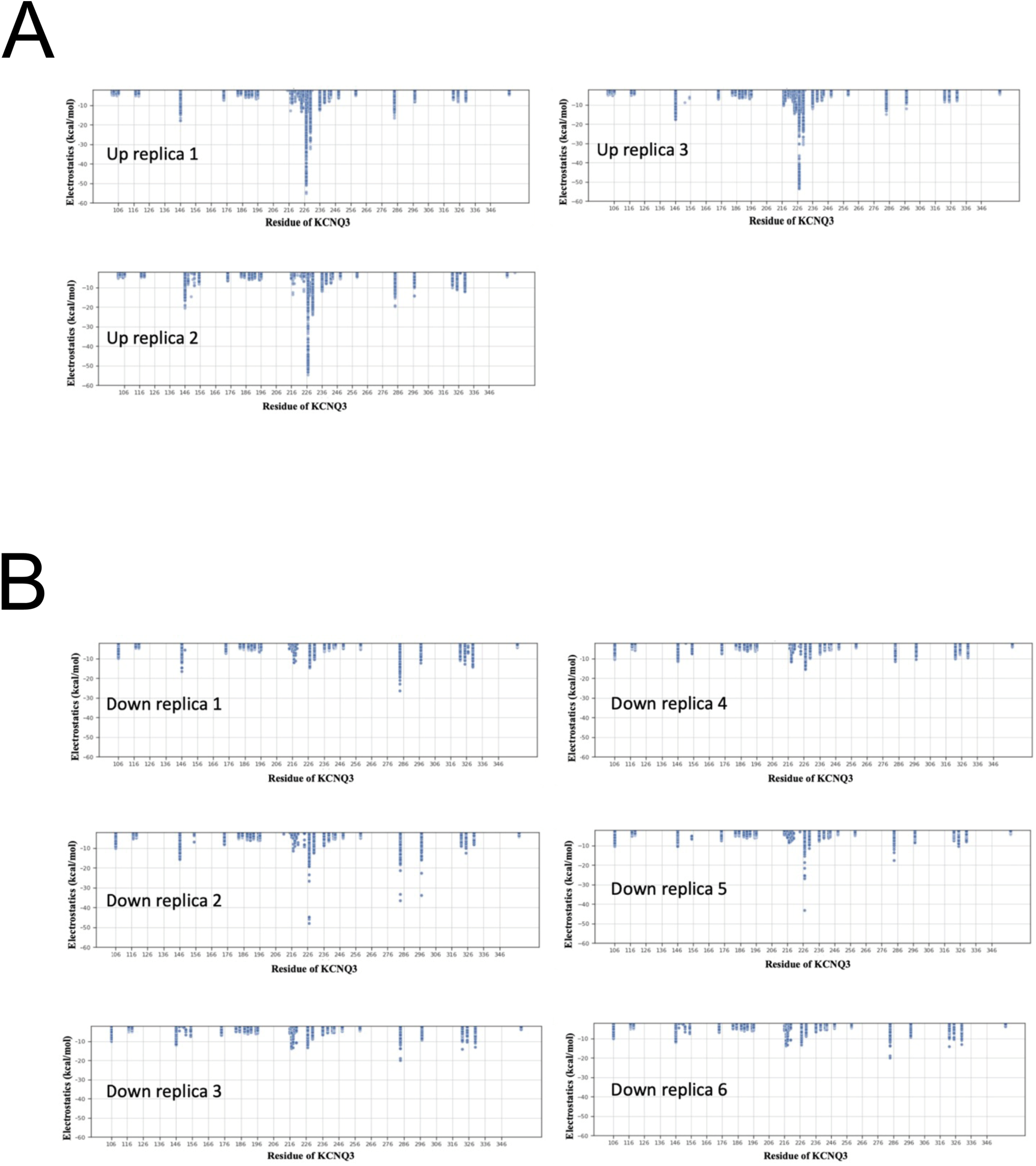
Molecular dynamics simulation predicts weaker NAT interaction with residues K146, R227, and K284 in the closed/resting state of KCNQ3. (A-B) Pairwise electrostatic interaction between NAT and KCNQ3 (A) active (S4 up) and (B) resting (S4 down) states. 3 replicas of S4 up state (A) and 6 replicas of S4 down state (B) trajectories are used for analysis. Each replica is 600 ns with frame size 6 ns. Each data point represents the electrostatic interaction from one of 100 snapshots over 600 ns. Only the favorable interactions < –2 kcal/mol were shown for clarity.

**Supplementary Figure 6.**
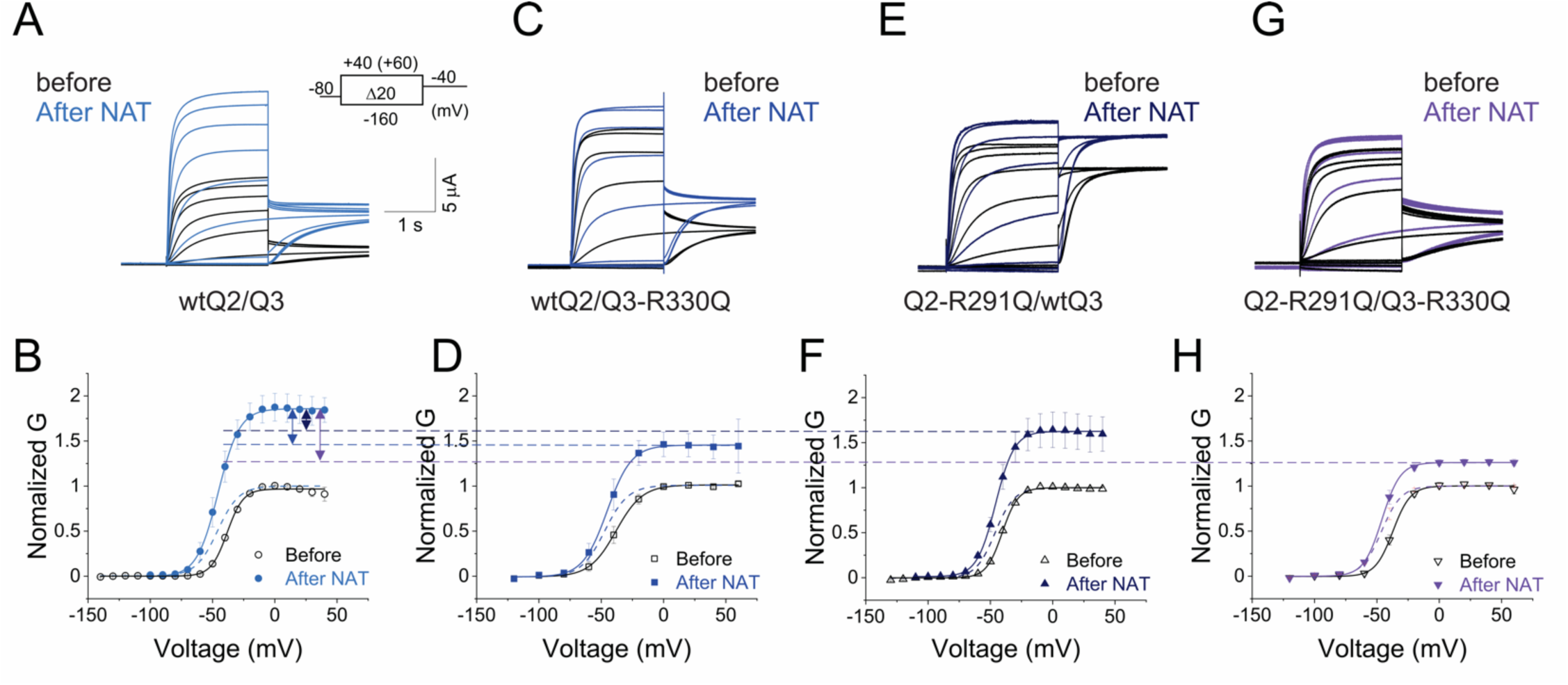
Residues R330 in the S6 helix of KCNQ3 is important for the Gmax effect of NAT. (*A*, *C, E,* and *G*) Representative current traces from (*A*) wild-type KCNQ2/3 (wtQ2/Q3) channels, (*C*) wt KCNQ2/KCNQ3-R330Q (wtQ2/Q3-R330Q), (*E*) KCNQ2-R291Q/wtKCNQ3 (Q2-R291Q/wtQ3), and (*G*) KCNQ2-R291Q/KCNQ3-R330Q (Q2-/R291Q/Q3-R330Q) channels in the absence (before) or presence (after) of 25 μM NAT (C) for the indicated voltage protocol. (*B*, *D, F,* and *H*) Steady-state conductance/voltage relationships, G(V)s, (solid lines from a Boltzmann fit) obtained from recordings of panels *A*, *C, E,* and G normalized to peak conductance before NAT application (open symbols). G(V) relationships (solid lines from a Boltzmann fit) in the presence of NAT are shown as closed symbols and dashed lines represent their respective normalized G(V). Data represents the mean ± SEM; n = 5–8. Values are given in Suppl. Table 2. Wild-type– (wtQ2/Q3) and mutant-bearing– heteromeric channels (wtQ2/Q3-*R330Q*, Q2-R291Q/wtQ3, and Q2-*R291Q/*Q3-*R330Q*) are expressed in a 1:1 ratio.

**Supplementary Figure 7.**
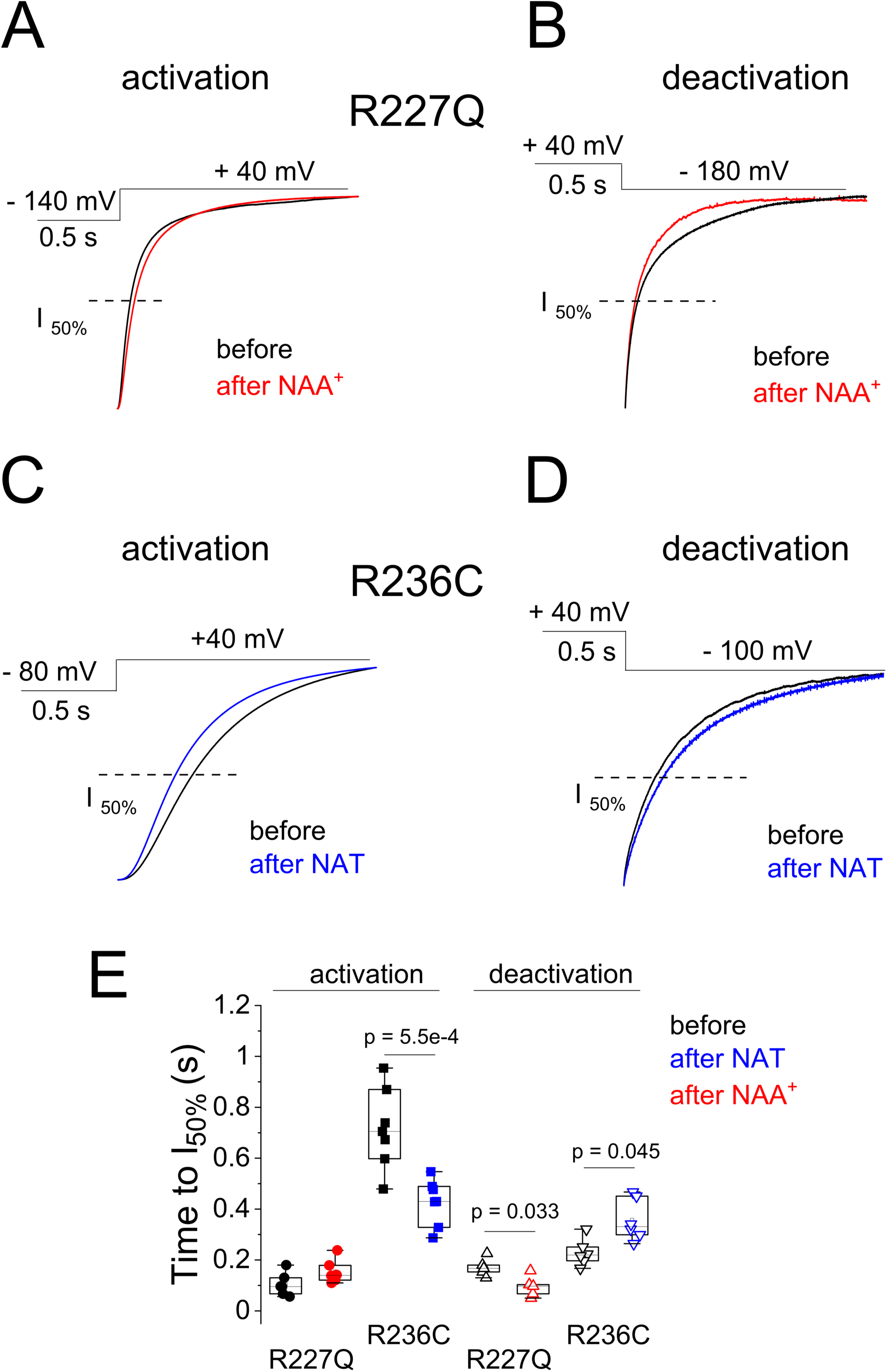
Effect of PUFAs on current activation and deactivation of NDD-associated mutations-R227Q and R236C. (*A*-*D*) Representative time courses of current activation (*A*, *C*) and deactivation (B, D) from KCNQ3-R227Q (*A*, *B*) and KCNQ3-R236C (*C*, *D*) channels in the absence (before) or presence (after) of 25 μM of NAA^+^ (red) or NAT (blue) in response to the indicated voltage steps. Dashed lines represent 50% maximum current level at the end of the depolarizing pulse. (*E*) Time courses of current activation (closed symbols) and deactivation (open symbols) in the absence (black) and presence of NAA^+^ (red) and NAT (blue) quantified as time to reach half maximum current level at the end of the depolarizing pulse in (A-D, dashed lines). Statistical significance determined using Student’s T-test to compare the PUFA–induced change in time to I_50%_. p < 0.05; only significant differences are shown.

**Supplementary Figure 8.**
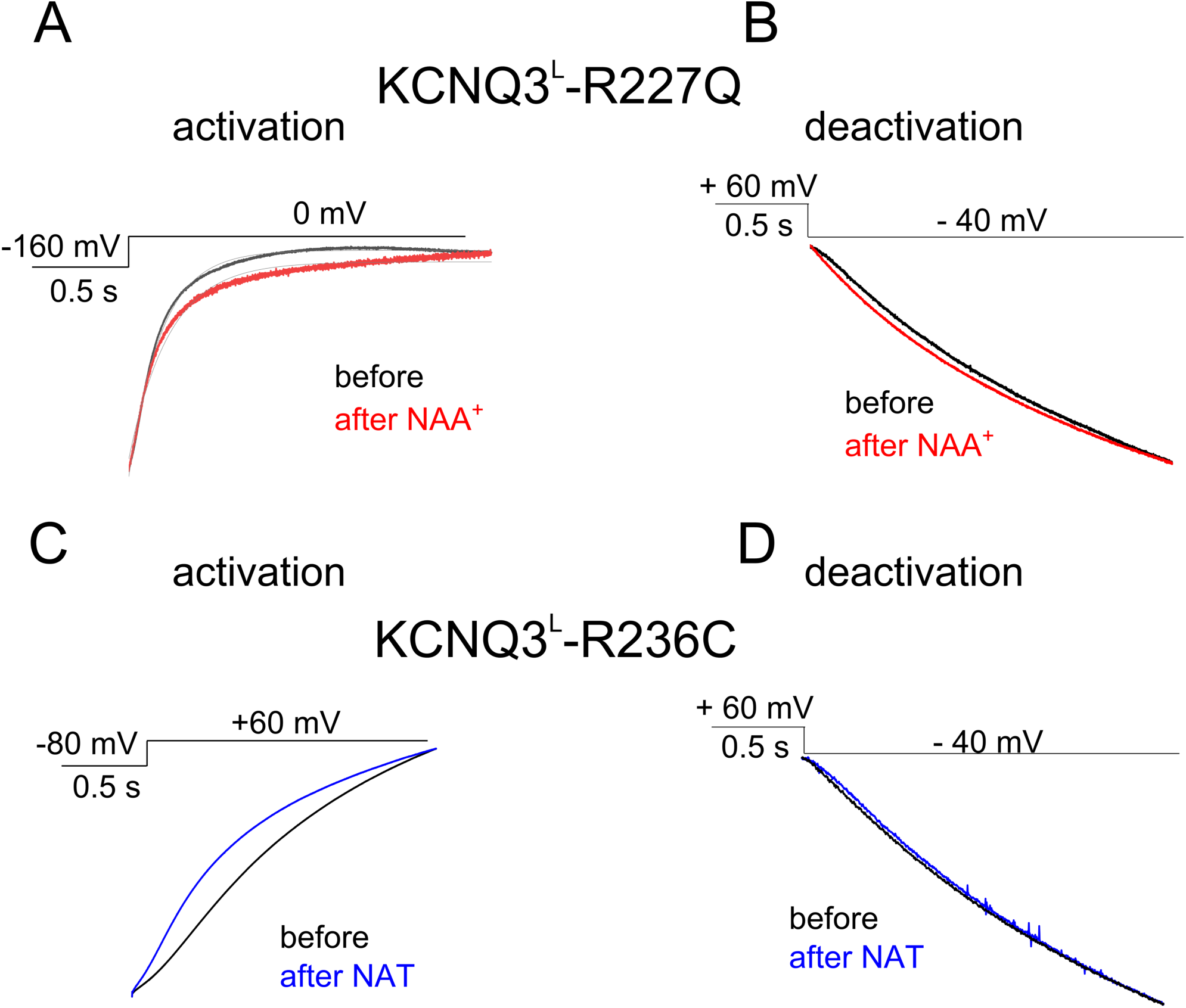
Effect of PUFAs on current activation and deactivation of labeled R1227Q and R236C channels. (A, B) Representative time courses of current activation (A, C) and deactivation (B, D) from Alexa488-labeled– KCNQ3^L^-R227Q (*A, B*) and KCNQ3^L^-R236C (*C, D*) channels in the absence (before) or presence (after) of 25 μM of NAA^+^ (red) or NAT (blue) in response to the indicated voltage steps.

**Supplemental Table 1.**
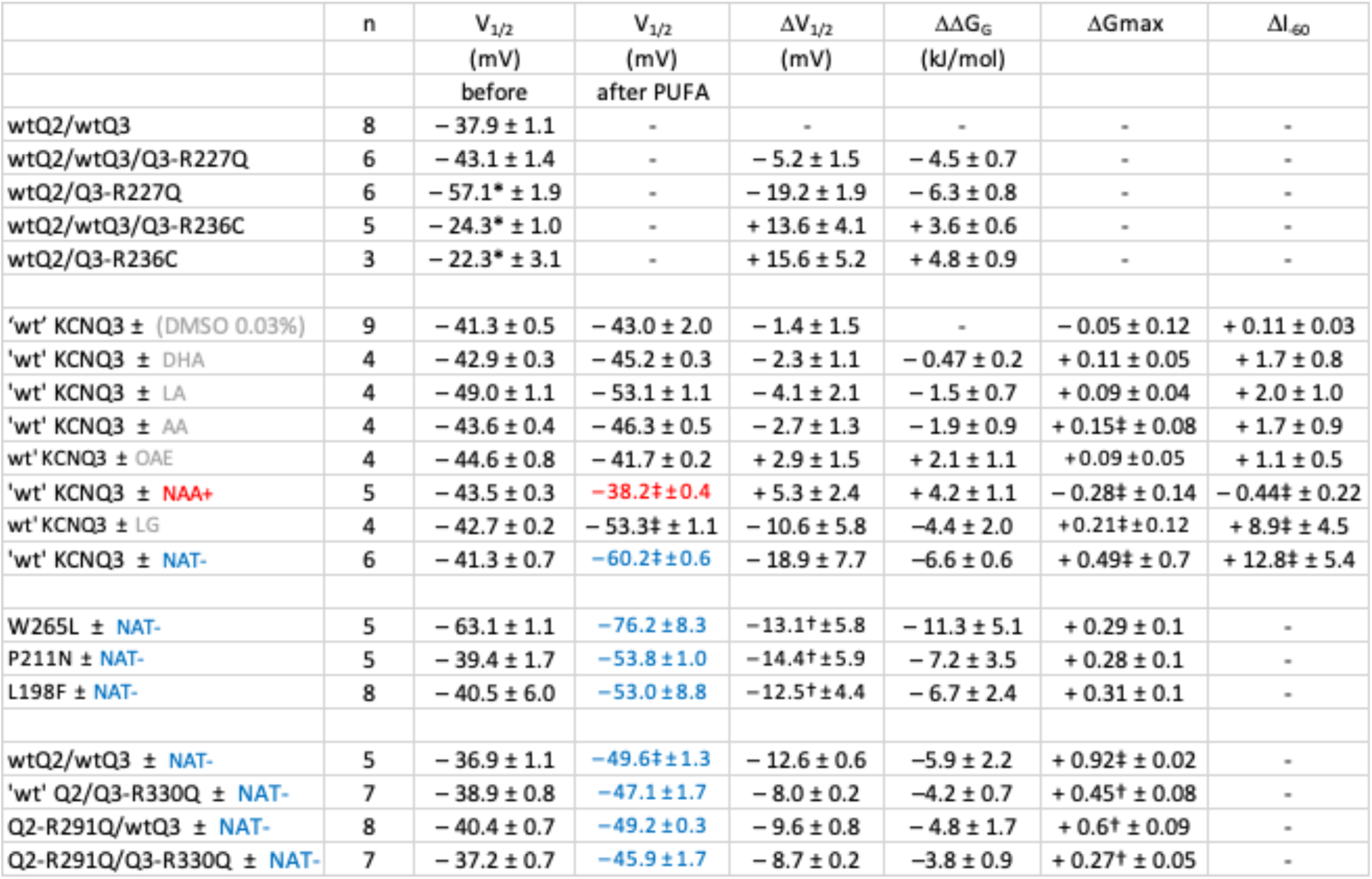
Effect of PUFAs and NDD-associated mutations on heteromeric KCNQ2/3 channels. Summary data for half-activation voltage (V_1/2_ and F_1/2_) for each G(V), shifts in half-activation voltage (ΔV_1/2_), Gibbs free energy associated with channel opening (11G_G_), maximal current amplitude (ΔGmax), and relative change in potassium current at –60 mV (1I_–60_) in the absence or presence of 25 μM PUFAs (color coded) on homomeric– and heteromeric wt (KCNQ3 or KCNQ2/3) and NDD-associated mutations. 5,8,11,14-cis-N-arachidonoyl amine (NAA^+^), 5,8,11,14-all-cis-N-arachidonoyl taurine (NAT) Values are means ± S.E. n represents the number of cells analyzed. Statistical significance determined using Student’s T-test or one-way ANOVA and Bonferroni’s post hoc test to compare the PUFA–induced change in ΔV_1/2_, ΔGmax, 11G, 1I_–60_, and time to I_50%_. p < 0.05 vs. the corresponding value in: *wt heteromeric KCNQ2/3, †wt homomeric or heteromeric KCNQ2/3 ± PUFA, and ‡wt homomeric KCNQ3 or heteromeric KCNQ2/3 ± DMSO

